# The *Hansenula polymorpha* Mitochondrial Carrier Family proteins Mir1 and Aac2 are dually localized at peroxisomes and mitochondria

**DOI:** 10.1101/2023.11.15.567167

**Authors:** Marc Pilegaard Pedersen, Justina C. Wolters, Rinse de Boer, Arjen M. Krikken, Ida J. van der Klei

## Abstract

Peroxisomes are ubiquitous cell organelles involved in various metabolic pathways. In order to properly function, several cofactors, substrates and products of peroxisomal enzymes need to pass the organellar membrane. So far only a few transporter proteins have been identified. We analysed peroxisomal membrane fractions purified from the yeast *Hansenula polymorpha* by untargeted label-free quantitation mass spectrometry. As expected, several known peroxisome-associated proteins were enriched in the peroxisomal membrane fraction. In addition, several other proteins were enriched, including mitochondrial transport proteins. Localization studies revealed that two of them, the mitochondrial carrier family proteins Aac2 and Mir1, have a dual localization on mitochondria and peroxisomes. To better understand the molecular mechanisms of dual sorting, we tested the localization of Mir1 in cells lacking Pex3 or Pex19, two peroxins that play a role in targeting of peroxisomal membrane proteins. In these cells Mir1 only localized to mitochondria, indicating that Pex3 and Pex19 are required to sort Mir1 to peroxisomes. Analysis of the localization of various truncated versions of Mir1 in wild-type *H. polymorpha* cells revealed that several localized to mitochondria, but only one, consisting of the transmembrane domains 3-6, was peroxisomal. Peroxisomal localization of this construct was lost in a *MIR1* deletion strain, indicating that full length Mir1 was required for the localization of the truncated protein to peroxisomes. Our data suggest that only full length Mir1 sorts to peroxisomes, while Mir1 contains multiple regions with mitochondrial sorting information.

## Introduction

Peroxisomes are small, single membrane bound organelles present in almost all eukaryotic cells. Mutations causing defects in peroxisome biogenesis or peroxisomal enzymes can cause severe life-threatening diseases in man, illustrating the importance of peroxisomes .

The peroxisomal matrix has the highest protein concentration of all cell organelles and consists of densely packed enzyme molecules that catalyse a plethora of anabolic and catabolic processes, such as the beta-oxidation of fatty acids, catabolism of (poly)amines, neutralization of reactive oxygen species and biosynthesis of several components (Wanders et al., 2023) (van der Klei and Veenhuis, 2013). Degradation and biosynthesis processes in the peroxisomal matrix imply that many molecules (enzyme co-factors, substrates, products) should pass the peroxisomal membrane. So far, only a few peroxisomal transporters have been identified. In addition, evidence has been presented that the peroxisomal membrane contains proteins that can form pores for small molecules (Chornyi et al., 2020). Moreover, peroxisomal matrix protein import involves the formation of (transient) pores, which may also allow diffusion of small molecules (Feng et al., 2022; Gao et al., 2022).

Identified mammalian peroxisomal transporters include ABCD transporters and monocarboxylate transporters (Chornyi et al., 2020). In *Saccharomyces cerevisiae* ABC transporters (Pxa1/Pxa2), adenine nucleotide transporters (Ant1, Aac2), and the oligo peptide transport Opt2 localize to peroxisomes (Elbaz-Alon et al., 2014; Plett et al., 2020; van Roermund et al., 2021). Recently, Pmp47 of the yeast *Debaryomyces hansenii* was shown to function also as peroxisomal NAD^+^/NADH exchanger (Turkolmez et al., 2023).

The mammalian peroxisomal membrane protein (PMP) Pxmp2 was the first protein suggested to function as a pore, which enables diffusion of molecules of up 300 Dalton (Rokka et al., 2009). A homologue of mammalian Pxmp2 is absent in yeast, but in *S. cerevisiae* the peroxin Pex11 was demonstrated to have pore-forming properties (Mindthoff et al., 2016). This observation is in line with data presented by Deloache and colleagues, who showed that *S. cerevisiae* peroxisomal membranes are permeable to some but not all small hydrophilic metabolites *in vivo* ((DeLoache et al., 2016).

Based on the above observations, different models exist for solute transfer across the peroxisomal membrane: 1) pore-forming proteins allow free and unhindered diffusion of small molecules up to approx. 300 Da, 2) selective transporters are responsible for transport of specific molecules or 3) both models are correct where pores enable diffusion of small molecules, while transporters are required for selective transport of larger ones.

Here we aimed to identify novel PMPs with transport functions by proteomics analysis of isolated peroxisomal membranes from the yeast *Hansenula polymorpha* (also called *Ogataea parapolymorpha, Ogataea polymorpha or Pichia angusta*). Whole organelle proteomics has been successfully applied with peroxisomes isolated from the yeasts *S. cerevisiae* (Marelli et al., 2004) and *Pichia pastoris* ((Valli et al., 2020). However, these analyses did not result in the identification of novel transport proteins. We therefore now used enriched peroxisomal membrane fractions from *H. polymorpha* in search for novel transport proteins. In *H. polymorpha* peroxisomes are essential for the metabolism of methanol. In carbon limited chemostat cultures peroxisomes are massively induced and may take up 80 % if the cell volume (Veenhuis et al., 1978). These cells are therefore ideal starting material to enrich peroxisomal membranes. We show that in addition to many known PMPs, mitochondrial porin and transporters of the mitochondrial carrier family (MCF) were enriched in the peroxisomal membrane fractions. Co-localization studies demonstrated that two MCF proteins, Aac2 and Mir1, showed a dual localization on peroxisomes and mitochondria. MCF proteins typically localize to the mitochondrial inner membrane (MIM). The sorting pathway of MCF proteins to the MIM has been studied extensively and proteins involved in transport across the mitochondrial outer membrane and insertion in the MIM are well characterized (Ferramosca and Zara, 2013). We here show that sorting of Mir1 to peroxisomes requires Pex3 and Pex19, two peroxins known to be involved in PMP sorting. Furthermore, we analysed the localization of several truncated Mir1 variants to identify regions with mitochondrial and peroxisomal sorting information. This revealed that only full length Mir1 sorts to peroxisomes.

## Results

### Isolation of peroxisomal membrane fractions

To identify novel peroxisomal transporters, *H. polymorpha* wild type (WT) cells were cultivated in a glucose/methanol limited chemostat to induce massive peroxisome proliferation. Peroxisomes were isolated from a post nuclear supernatant (PNS) by sucrose density gradient centrifugation (Fig. 1A). The sucrose gradients were fractionated from the bottom and fractions were subjected to immunoblot analysis using anti-Pex11 antibodies to identify fractions enriched with peroxisomes (Fig. 1B and Fig. S1). Mitochondria are generally the primary contaminants upon peroxisome isolation procedures. Because antibodies against *H. polymorpha* mitochondrial proteins are not available, we introduced GFP N-terminally tagged with the mitochondrial presequence of *N. crassa* subunit 9 of the F_0_ATPase (su9-GFP) (Westermann and Neupert, 2000) and used anti-GFP antibodies to detect mitochondria by Western blotting (Fig. 1B). A representative gradient and the corresponding Western blots are shown in Fig. 1B.

**Figure. 1:**
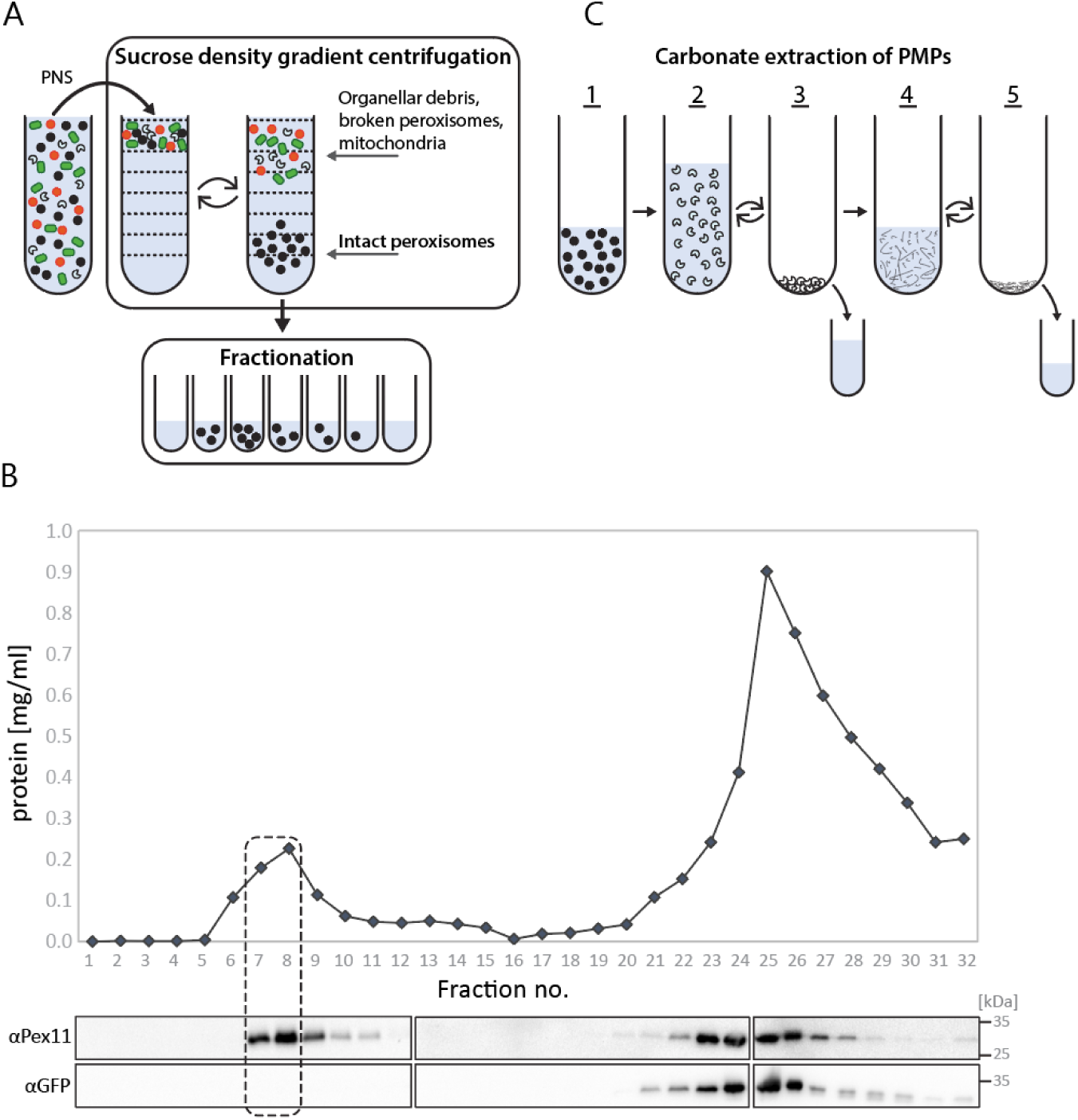
Peroxisome isolation from *H. polymorpha* producing su9-GFP grown in a glucose/methanol-limited chemostat. **A)** Schematic representation of sucrose density centrifugation of a PNS and collection of gradient fractions. **B)** Graph showing protein profile of a representative sucrose gradient and western blot analysis of all fractions using anti-Pex11 (peroxisomes) or anti-GFP (to detect the mitochondrial marker su9-GFP) antibodies. Equal volumes were loaded per lane. Fractions 7 and 8 were pooled for further analysis. **C**): Schematic representation of the isolation of the peroxisomal membrane fraction. 1) pooling of peroxisomal fractions, 2) osmotic lysis of peroxisomal fractions followed by ultra-centrifugation, 3) removal of supernatant, 4) carbonate extraction of pellet fraction followed by ultra-centrifugation, 5) removal of supernatant. The PNS and membrane pellet after carbonate extraction (5) were subjected to MS/MS proteomic analysis.

Immunoblotting using anti-Pex11 antibodies revealed that Pex11 is present in the high sucrose density region (fractions 7-9), but also in the lower sucrose density region (fractions 20-29). The latter may be due to unbroken protoplasts and/or broken peroxisomes present in the PNS. Immunoblotting with anti-GFP antibodies revealed that the mitochondrial marker only occurred in the lower sucrose density region (fraction 23-27), as expected. Only upon a 20 x longer exposure time a very faint GFP band could be detected in the high sucrose density region as well (Fig. S1). These results shown that only very minor mitochondrial contamination occurred in the peroxisomal peak fractions. Because we aimed to identify novel peroxisomal transporters, which are integral membrane proteins, we subjected the peroxisomal peak fractions to osmotic lysis and carbonate extraction, to remove soluble and membrane associated proteins (Fig. 1C). For each of the five biological replicates, the peroxisomal peak fractions were pooled and osmotically lysed. After ultracentrifugation the supernatant was separated from the pellet, which was subjected to carbonate extraction. The pellet obtained after ultracentrifugation was used for mass spectrometry (Fig. 1C and Table S1) and designated PMEM (peroxisomal membrane fraction).

### Mass spectrometry analysis of the PNS and PMEM

To identify novel peroxisomal transporters, we performed untargeted label-free quantitation (LFQ) mass spectrometry (MS) on PMEM samples obtained from 5 biological replicates. A similar analysis was performed on the corresponding PNS fractions to monitor which membrane proteins enriched in PMEM (Table 1 and Suppl. Tables S1 and S2). Proteins were ranked based on the signal intensities detected in the mass spec and the ranking was used to identify the proteins that were relatively enriched in the PMEM fraction compared to the PNS fraction.

**Table 1:** Protein composition of 40 highest ranked proteins in PNS and PMEM. The relative protein signal intensities ranking compares the median corrected Log2-LFQ values of ranked proteins from **A)** PNS or **B)** PMEM. The median corrected Log2-LFQ values are an average of five independent biological replicates (n=5). The top 40 proteins for both samples are shown. The complete tables can be found in Suppl. Tables S1 and S2. Proteins marked in bold: PMPs. Proteins underlined mitochondrial carriers or pore forming proteins. UniProt entry name for each listed open reading frame (ORF) and protein descriptions were extracted from the *Ogataea parapolymorpha* proteome (proteome ID #UP000008673). *H. polymorpha* proteins that have not been described before were named after their closest homolog in *S. cerevisiae* or *P. pastoris*. Protein localization: Px: peroxisome, Mi: mitochondria, Cy: cytosol, Pm: plasma membrane, LD: lipid droplet, Va: vacuole, ER: endoplasmic reticulum, Nu: nucleus, N/A: unknown localization, - proteins which have not yet been named. Due to the high sequence identity of the malate dehydrogenases *(Mdh1, Mdh2), transaldolases **(Tal1, Tal2), and long chain fatty acyl-CoA synthases ***(Faa1, Faa2, Faa3, Faa4) it was not possible to distinguish between their homologs.

| PNS |  |  |  |  | PMEM |  |  |  |  |
| --- | --- | --- | --- | --- | --- | --- | --- | --- | --- |
| ORF | Name | Protein description | Localization | Median corrected Log2-LFQ | ORF | Name | Protein description | Localization | Median corrected Log2-LFQ |
| W1QCJ3 | Aox | Alcohol oxidase | Px | 38,05 | W1QIW6 | Das | Dihydroxyacetone synthase | Px | 32,66 |
| W1QIW6 | Das | Dihydroxyacetone synthase | Px | 35,38 | W1QFE1 | Pex11 | **Pex11** | Px | 32,43 |
| W1QJ02 | Cta | Catalase | Px | 34,69 | W1QCJ3 | Aox | Alcohol oxidase | Px | 32,36 |
| W1Q6W1 | Atp1 | ATP synthase subunit alpha | Mi | 33,22 | W1QJ22 | Por1 | <u>Mitochondrial outer membrane protein porin</u> | Mi | 30,39 |
| W1QA59 | Atp2 | ATP synthase subunit beta | Mi | 33,16 | W1QES8 | Pmp20 | Putative peroxiredoxin-B | Px | 30,23 |
| W1QFP1 | Adh2 | Alcohol dehydrogenase 2 | Cy | 32,96 | E7R3W5 | - | Histone H4 | Nu | 29,96 |
| W1QJ22 | Por1 | <u>Mitochondrial outer membrane protein porin</u> | Mi | 32,86 | E7R2Y5 | Eft1 | Elongation factor 1 | Px, Cy | 29,25 |
| W1QBL6 | Aat1 | Aspartate aminotransferase | Px | 32,73 | W1Q7K8 | Pcs60 | Peroxisomal-coenzyme A synthetase (Fat2) | Px | 29,16 |
| E7R2Y5 | Eft1 | Elongation factor 1 | Px, Cy | 32,68 | W1QJ02 | Cta | Catalase | Px | 29,10 |
| W1QES8 | Pmp20 | Putative peroxiredoxin-B | Px | 32,49 | W1Q6U7 | Aac2 | <u>ADP/ATP carrier protein</u> | Mi | 28,99 |
| W1Q6V1 | Fld1 | S-(hydroxymethyl)glutathione dehydrogenase | Cy | 32,34 | W1QF22 | Pma1 | Plasma membrane ATPase | Pm | 28,72 |
| W1QGV6 | Mdh* | Malate dehydrogenase | Mi, Px, Cy | 32,22 | W1QG51 | Pmp47 | **PMP47** | Px | 27,78 |
| O59841 | Tdh1 | Glyceraldehyde-3-phosphate dehydrogenase | Cy | 32,16 | W1QFP1 | Adh2 | Alcohol dehydrogenase 2 | Cy | 26,97 |
| W1QKD3 | Tal** | Transaldolase | Px, Cy | 31,66 | O74258 | Act | Actin | Cy, Px | 26,80 |
| W1QK02 | Mdh* | Malate dehydrogenase | Mi, Px, Cy | 31,62 | W1QLS6 | Ubi4 | Ubiquitin-40S ribosomal protein | N/A | 26,20 |
| W1QFE1 | Pex11 | **Pex11** | Px | 31,60 | W1QA59 | Atp2 | ATP synthase subunit beta | Mi | 25,90 |
| W1Q6U7 | Aac2 | ADP/ATP carrier protein | Mi | 31,59 | W1QFF2 | Pex2 | **Pex2** | Px | 25,61 |
| W1QH90 | Met6 | Homocysteine S-methyltransferase | Cy, Pm | 31,58 | W1QHN4 | Vac8 | Vacuolar protein 8 | Va | 24,99 |
| W1QKD7 | Tal** | Transaldolase | Px, Cy | 31,47 | W1Q838 | Tpi1 | Triosephosphate isomerase | Cy, Mi, Pm | 24,51 |
| O74258 | Act | Actin | Cy, Px | 31,44 | W1QB82 | - | SAP domain-containing protein | N/A | 23,03 |
| W1QEC3 | Fba | Fructose-bisphosphate aldolase | Cy | 31,44 | W1QEN7 | Snf3 | High-affinity glucose transporter SNF3 | Pm, ER | 22,75 |
| W1QD17 | Hsp60 | Heat shock protein 60 | Mi | 31,42 | W1QH28 | - | Histone H2A | Nu | 22,38 |
| W1Q801 | Fdh1 | Formate dehydrogenase | Cy | 31,40 | W1QKD7 | Tal** | Transaldolase | Px, Cy | 22,24 |
| W1Q838 | Tpi1 | Triosephosphate isomerase | Cy, Mi, Pm | 31,30 | W1Q7W6 | Pho88 | Functions in the SND pathway PHO88 | ER, Mi | 22,08 |
| W1Q9M7 | Acs1 | Acetyl-coenzyme A synthetase | Cy, Mi, Nu | 31,30 | W1QL03 | Mir1 | <u>Mitochondrial phosphate carrier protein</u> | Mi | 21,99 |
| W1QBG5 | Hsp90 | ATP-dependent molecular chaperone HSC82 | Cy | 31,22 | W1Q8B2 | Odc1 | <u>Mitochondrial 2-oxodicarboxylate carrier 2</u> | Mi | 21,92 |
| W1QD98 | - | S-formylglutathione hydrolase | Cy | 31,17 | W1QHx5 | Yet3 | ER transmembrane protein | ER | 21,83 |
| W1QF08 | Dak | Dihydroxyacetone kinase | Cy | 31,13 | W1QGW3 | Vph1 | V-type proton ATPase subunit a | Va | 21,80 |
| W1Q8A9 | Aat2 | Aspartate aminotransferase | Px | 31,11 | W1QKD3 | Tal** | Transaldolase | Px, Cy | 21,72 |
| W1QIU6 | Eno1 | Phosphopyruvate hydratase | Cy | 30,91 | W1Q9K4 | Pex12 | **Pex12** | Px | 21,70 |
| W1Q7H2 | Eft2 | Elongation factor 2 | Cy | 30,82 | W1Q6W1 | Atp1 | ATP synthase subunit alpha | Mi | 21,54 |
| W1Q894 | Ssc1 | Heat shock protein SSC1 | Mi | 30,72 | W1Q961 | Rki1 | D-ribose-5-phosphate ketol-isomerase | Cy | 21,38 |
| W1QIC9 | Pgk1 | Phosphoglycerate kinase | Cy, Mi | 30,69 | W1QDA8 | Rho1 | GTP-binding protein RHO1 | Px, Pm | 21,24 |
| W1QLK4 | Aco1 | Aconitate hydratase | Mi | 30,60 | W1Q8E4 | Faa*** | Long chain fatty acyl-CoA synthetase | Cy, Pm, LD | 21,12 |
| W1QKK3 | Gnd | 6-phosphogluconate dehydrogenase | Cy | 30,57 | W1QJ84 | Pex10 | **Pex10** | Px | 21,07 |
| W1Q8A6 | Ilv5 | Acetohydroxy-acid reductoisomerase | Mi | 30,56 | W1QL75 | Opi3 | Methyltransferase | ER, Mi | 21,06 |
| W1QEM2 | - | Ubiquinol cytochrome-c reductase core protein 2 | N/A | 30,40 | W1QFR9 | Atg37 | **Autophagy-related protein 37** | Px | 20,82 |
| W1QLI1 | Fas1 | Fatty acid synthase subunit beta | Cy, Mi, LD | 30,36 | W1QA67 | Tsc13 | Enoyl reductase | ER | 20,68 |
| W1QF22 | Pma1 | Plasma membrane ATPase | Pm | 30,33 | W1QFG9 | Pex3 | **Pex3** | Px | 20,07 |
| W1QHM6 | Ino1 | Inositol-3-phosphate synthase | Cy | 30,25 | W1QBL6 | Aat1 | Aspartate aminotransferase | Px | 19,01 |

Yeast PNS fractions commonly contain cell organelles (peroxisomes, mitochondria, ER, etc.) and the cytosol (Fig. 1A). Indeed, predominantly proteins of these cellular compartments were detected in the PNS (Table 1, Suppl Table S2). The top 3 hits of the PNS are the peroxisomal matrix enzymes alcohol oxidase (Aox), dihydroxyacetone synthase (Das) and catalase (Cta), which is in line with the high abundance of these proteins in methanol containing carbon-limited chemostat cultures (van der Klei et al., 2006). Pex11, an abundant PMP in yeast (Marshall et al., 1995), is also ranked relatively high in the PNS. Most of the other top ranked PNS proteins are housekeeping proteins and/or from cytosolic metabolic pathways such as the glycolysis and the oxidative -and non-oxidative pentose phosphate pathways (Table 1). As expected, in PMEM much less housekeeping proteins and cytosolic enzymes were detected (Suppl. Table S1). Instead, many known peroxisomal or peroxisome associated proteins are present in the top ranked PMEM proteins. Because of the high number of peroxisomal proteins in the top 40 ranked PMEM proteins, we confine to these ones for more detailed descriptions (Table 1).

The top 40 ranked PMEM proteins contained 7 known PMPs, while there was only one, Pex11, in the top 40 ranked PNS proteins. In the PMEM Pex11 is the second highest ranked protein, whereas it was ranked 16 in the PNS (Table 1). Other known PMPs in the top 40 PMEM proteins include Pex3 (receptor involved in PMP sorting), Pex2, Pex10 and Pex12 (RING complex), Pmp47 (peroxisomal ATP:ADP/AMP exchanger), and Atg37 (the homologue of the *P. pastoris* autophagy protein Atg37 (Nazarko, 2014)).

The peroxisomal enzymes Das, Aox and Cta were also among the top 40 PMEM proteins, indicating that these matrix proteins were not completely separated from the peroxisomal membranes upon carbonate extraction. Similarly, the peroxisomal matrix enzymes transketolase (Tal2; (Kurylenko et al., 2018) and peroxiredoxin-B (called Pmp20 in *H. polymorpha* (Bener Aksam et al., 2008)) were detected in PMEM. Although the name of Pmp20 suggests that it is a membrane protein, Pmp20 is a soluble enzyme that associates to peroxisomal membranes. *S. cerevisiae* Pcs60 is also peroxisomal matrix enzyme that associates to peroxisomal membranes (Blobel and Erdmann, 1996). Similarly, we found the *H. polymorpha* homologue of *Sc*Pcs60 among the top 40 PMEM proteins. *S. cerevisiae* contains four long chain fatty acyl-CoA synthetases (Faa1-4), of which one is peroxisomal (Faa2). The PMEM also contains one Faa protein, but due to the high sequence identity, we could not establish whether this is the *Sc*Faa2 homolog. Notably, the *H. polymorpha* proteome only contain one protein described as a long chain fatty acyl-CoA synthetase (W1Q8E4).

Five additional proteins in the top 40 candidates of PMEM were previously shown to associate with yeast peroxisomes. These include *H. polymorpha* translation elongation factor 1-alpha (Kiel et al., 2007) and the vacuolar membrane protein Vac8 (Singh et al., 2020). The *H. polymorpha* homologs of *S. cerevisiae* actin and *Sc*Rho1, a GTPase involved in actin organization, were also among the top 40 PMEM proteins. Both proteins also associate with *S. cerevisiae* peroxisomes (Marelli et al., 2004). Recently the *S. cerevisiae* mitochondrial Aac2 protein was shown to have a dual localization at mitochondria and peroxisomes (van Roermund et al., 2021). The presence of its *H. polymorpha* homolog in the PMEM may therefore also be a result of dual localization of this protein in *H. polymorpha* mitochondria and peroxisomes.

Summarizing, the top 40 candidates of the PMEM prepared from *H. polymorpha* cells contains many proteins that were previously were indicated to localize yeast to peroxisomes. Moreover, these peroxisomal proteins invariably ranked higher in the PMEM relative to the PNS, confirming that peroxisomal proteins are highly enriched among the top 40 PMEM proteins.

The top 40 PMEM proteins contains one uncharacterized protein (W1QB82), which has an SAP domain that binds DNA/RNA. In addition the PMEM has many (membrane) proteins of other cell organelles, especially from mitochondria (Atp1, Atp2, Aac2, Mir1, Odc1, Por1). Four of them (Por1, Mir1, Aac2 and Odc1) were absent in the top 40 proteins of the PNS, indicating that they are co-enriched with peroxisomal proteins in the PMEM, while the remaining two (Atp1 and Atp2) were lower on the list of the PMEM proteins. Because the four co-enriched proteins all have a transport function, we next experimentally tested their localization.

### Aac2 and Mir1 have a dual localization on peroxisomes and mitochondria

To test whether the four mitochondrial membrane proteins, Por1, Mir1, Aac2 and Odc1, have a dual localization on peroxisomes and mitochondria in *H. polymorpha*, we performed confocal laser scanning microscopy (CLSM) with Airyscan imaging. To this end, we constructed strains producing C-terminal GFP fusions under control of relatively strong, constitutive promoters. Pex3-mKate2 under control of its endogenous promoter was introduced as peroxisomal marker. We imaged cells that were grown on media containing glucose or methanol. The advantage of imaging glucose-grown cells is that these cells generally contain one small peroxisome, while methanol grown cells are crowded with large peroxisomes making it more difficult to discriminate between peroxisomes and mitochondria.

Aac2-GFP exhibited tubular patterns indicative of mitochondrial localization. In addition, Aac2-GFP also was present on small puncta in glucose grown cells. These puncta represent peroxisomes because they also contain Pex3-mKate2 (Fig. 2A). Localization of Aac2-GFP on peroxisomes could not clearly be established in cells cultivated on methanol (Fig. 2B). Ocd1 also localized to mitochondria, however no Odc1-GFP signal was observed to colocalize with Pex3-mKate2 in glucose or methanol cultivated cells (Fig. 2 A, B). Cells producing Por1-GFP exhibited a punctate pattern, however these puncta did not contain Pex3-mKate2 and hence do not represent peroxisomes (Fig. 2A, B). Mir1-GFP showed both a mitochondrial and punctate pattern. The Mir1-GFP spots observed in glucose-grown cells colocalized with Pex3-mKate2, indicative for a dual localization of Mir1-GFP on peroxisomes and mitochondria. In methanol grown cells larger spherical structures also showed Mir1-GFP fluorescence in addition to mitochondria. These represent the larger peroxisomes that are present in methanol-grown cells, because they also contained Pex3-mKate2 (Fig. 2B). Control experiments confirmed that the Aac2-GFP and Mir1-GFP spots that co-localized with Pex3-mKate were not due to bleed through of Pex3-mKate2 in the GFP channel (Fig. 2A, B). We therefore conclude that Aac2 and Mir1 have a dual localization on mitochondria and peroxisomes.

**Figure 2:**
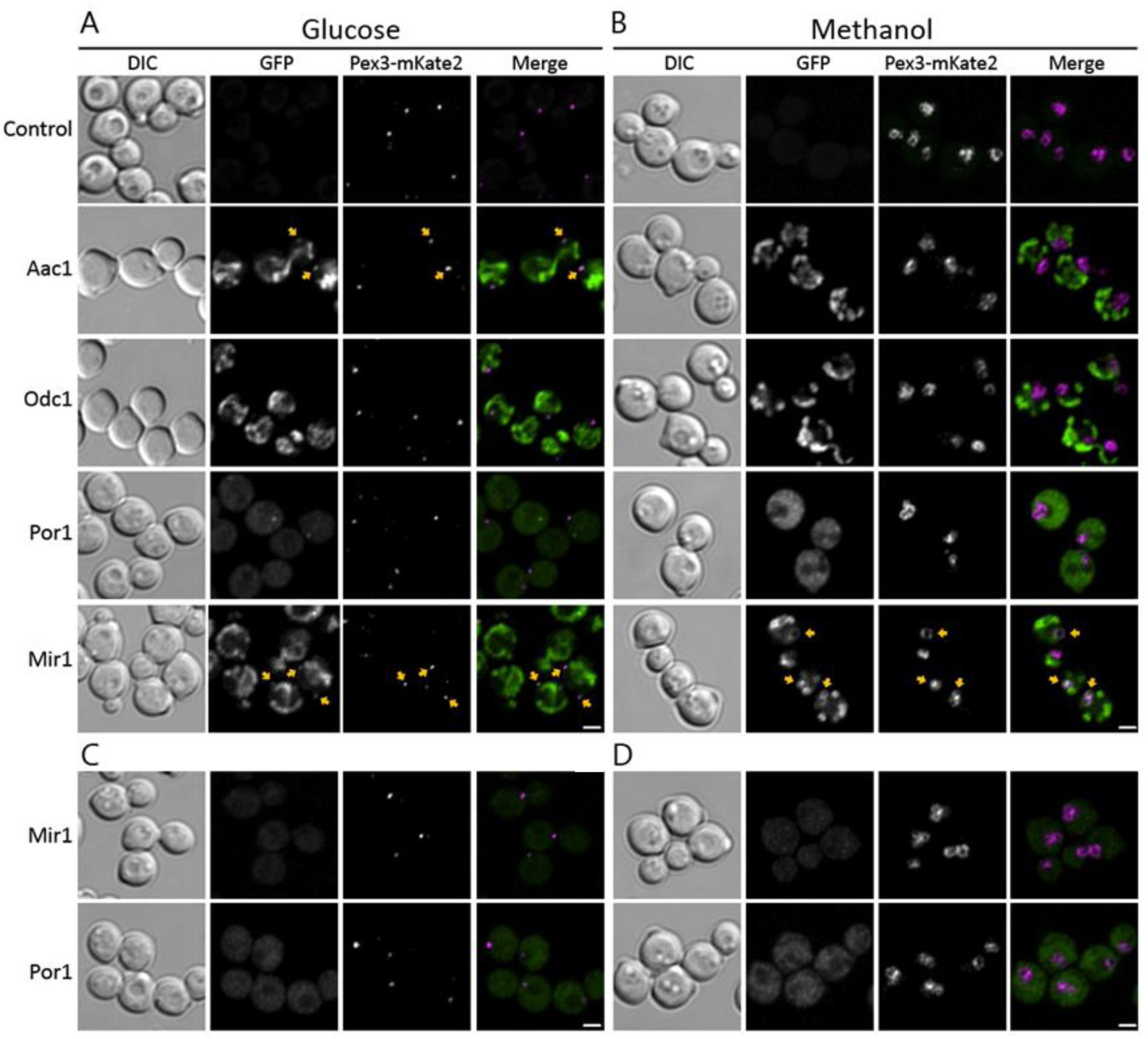
Aac2 and Mir1 have a dual localization on peroxisomes and mitochondria. Single focal plane CLSM Airyscan images of **A)** glucose-grown cells and **B)** methanol grown cells producing overexpressed C-terminal GFP fusions for the indicated proteins. **C)** glucose-grown cells and **D)** methanol grown cells producing overexpression N-terminal GFP fusions of GFP-Mir1 and GFP-Por1. Pex3-mKate2 was used as marker for all strains. The arrows indicate colocalization with Pex3-mKate2. Scale bar 1µm.

Because Por1-GFP was mislocalized, we also analysed a strain overproducing an N-terminal GFP fusions. However, this protein was also mislocalized to the cytosol.

Since peroxisomal Mir1-GFP localization is evident in both glucose and methanol-grown cells, we decided to further focus on the dual localization of Mir1. As shown in Fig. 2C, the localization of the N-terminal GFP fusion (GFP-Mir1) could not be established due to very low fluorescence signal (Fig. 2 C, D). We therefore continued our studies with the C-terminal fusion protein, Mir1 -GFP.

To exclude that the peroxisomal localization of Mir1-GFP is an artifact due to overexpression, we also expressed Mir1-GFP under control of its endogenous promoter. The Mir1-GFP fusion protein is functional, because cells of a *MIR1* deletion strain, but not cells producing endogenous Mir1-GFP, show a very severe growth defect on glucose media (Fig. S2). As shown in Fig. 3A, endogenously expressed Mir1-GFP colocalizes with mitochondria and Pex3-mKate2 spots, similar as observed for overexpressed Mir1-GFP. We also localized endogenously expressed Mir1-GFP in cells in which mitochondria were marked with the fluorescent dye MitoTracker. As shown in Fig. 3B, the bulk of the Mir1-GFP colocalizes with MitoTracker, confirming the mitochondrial localization of Mir1-GFP. In addition, Mir1-GFP localizes to puncta, which do not show MitoTracker fluorescence, indicating that Mir1-GFP indeed also localizes to other structures than mitochondria.

**Figure 3:**
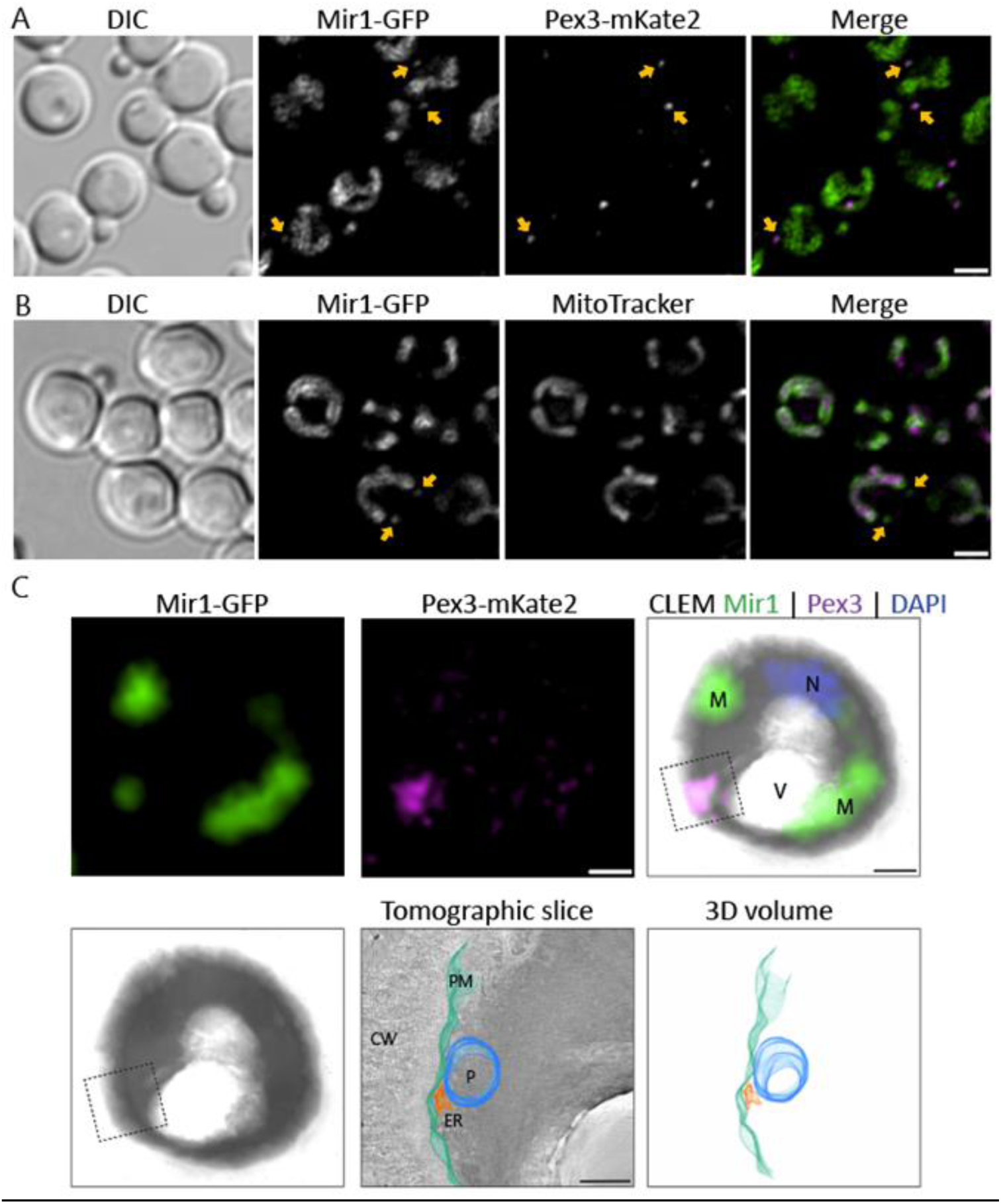
Mir1 localizes on mitochondria and peroxisomes. Single focal plane confocal laser scanning microscopy Airyscan images of glucose-grown cells expressing Mir1-GFP under control of the endogenous promoter together with **A)** Pex3-mKate2 as peroxisomal marker or **B)** MitoTracker™ Red to stain mitochondria. The arrows in **A** indicate colocalization of Mir1-GFP with Pex3-mKate2 and in **B** the position of the Mir1-GFP spot. Scalebars 1 µm. **C)** The upper row shows FM images and an overlay of FM and EM images of 150-nm thick cryosections of the same cell section. CLEM analysis of single focal plane of endogenously expressed Mir1-GFP localization and Pex3-mKate2 grown on glucose. Vacuole (V); mitochondria (M); nucleus (N). Lower row panels show reconstructed tomogram and 3D rendered. Peroxisome (P), blue; ER, orange; plasma membrane (PM), cyan; cell wall (CW). Scale bars in upper row 500 nm and in lower row (tomographic slice) 200 nm.

Finally, we performed correlative light and electron microscopy (CLEM) to analyse endogenous Mir1–GFP localization at high resolution using a strain also producing Pex3-mKate2. As expected, FM imaging revealed colocalization between Mir1-GFP and Pex3-mKate2 on separate punctate spots (Fig. 3C; upper panel). EM tomography shows that this spot represents a peroxisome. No mitochondria are present in the region where Mir1-GFP colocalizes with Pex3-mKate2 (Fig 3C; lower panel), indicating that indeed a portion of the Mir1-GFP protein localizes to peroxisomes.

### Peroxisomal sorting of Mir1 depends on Pex3 and Pex19

Pex3 and Pex19 are peroxins required for insertion of most PMPs into peroxisomal membranes. In the absence of these peroxins a minor subset of PMPs still localizes to peroxisomal membrane structures (e.g., Pex14), while other PMPs (especially those with multiple membrane spans) are mistargeted and/or degraded (Jansen and van der Klei, 2019). We analysed the localization of Mir1-GFP in Δ*pex3*Δ*atg1* and Δ*pex19*Δ*atg1* double deletion strains. *ATG1* was deleted in these strains because the Pex14-containing peroxisomal membrane structures are sensitive to autophagic degradation and only easily detectable upon blocking autophagy (Knoops et al., 2014). In both double deletion strains Mir1-GFP localizes to mitochondria but is absent on the peroxisomal membrane structures marked with Pex14-mCherry (Fig. 4). These data show that Mir1 targeting to peroxisomes depends on Pex3 and Pex19.

**Figure 4:**
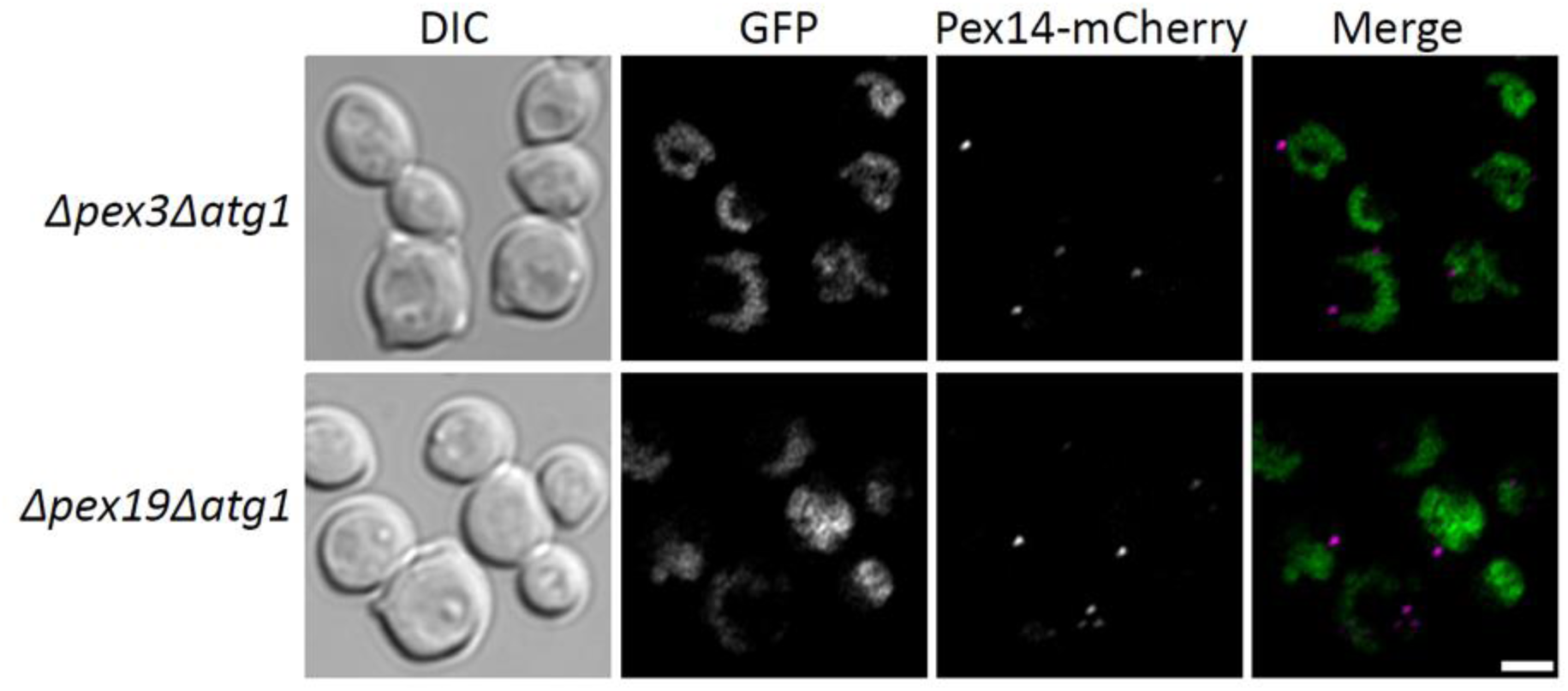
Peroxisomal localization of Mir1 depends on the presence of Pex3 and Pex19. Single focal plane confocal laser scanning microscopy Airyscan images of glucose-grown cells expressing C-terminal fusion of Mir1-GFP under control of the endogenous promoter. Images represent the Δ*pex3*Δ*atg1* and Δ*pex19*Δ*atg1* double mutant with Pex14-mCherry peroxisomal marker. Representative images of three independent experiments. Scale bar 1µm.

### The Mir1 homolog Pic2 does not localize to peroxisomes

In *S. cerevisiae*, Mir1 has a functionally redundant homolog, Pic2, which facilitates transport of copper in addition to phosphate across the MIM (Takabatake et al., 2001) (Vest et al., 2013). *H. polymorpha* also contains the Pic2 (W1QKL5) homolog of Mir1, which shows an amino acid sequence identity of 35% and a sequence similarity of 56% with *Hp*Mir1. Sequence alignment indicated that Pic2 has three gaps when compared to Mir1 (see Fig. 5A). Since Pic2 was not enriched in PMEM (Suppl. Table S1), we assumed that it would not be dually localized. To test this, Pic2 was C-terminally fused to GFP and expressed under control of its endogenous promoter. Consistent with the outcome of the proteomic analysis, Pic2-GFP displayed only tubular structures resembling mitochondria with no distinct punctate patterns co-localizing with Pex3-mKate2 (Fig. 5B).

**Figure 5:**
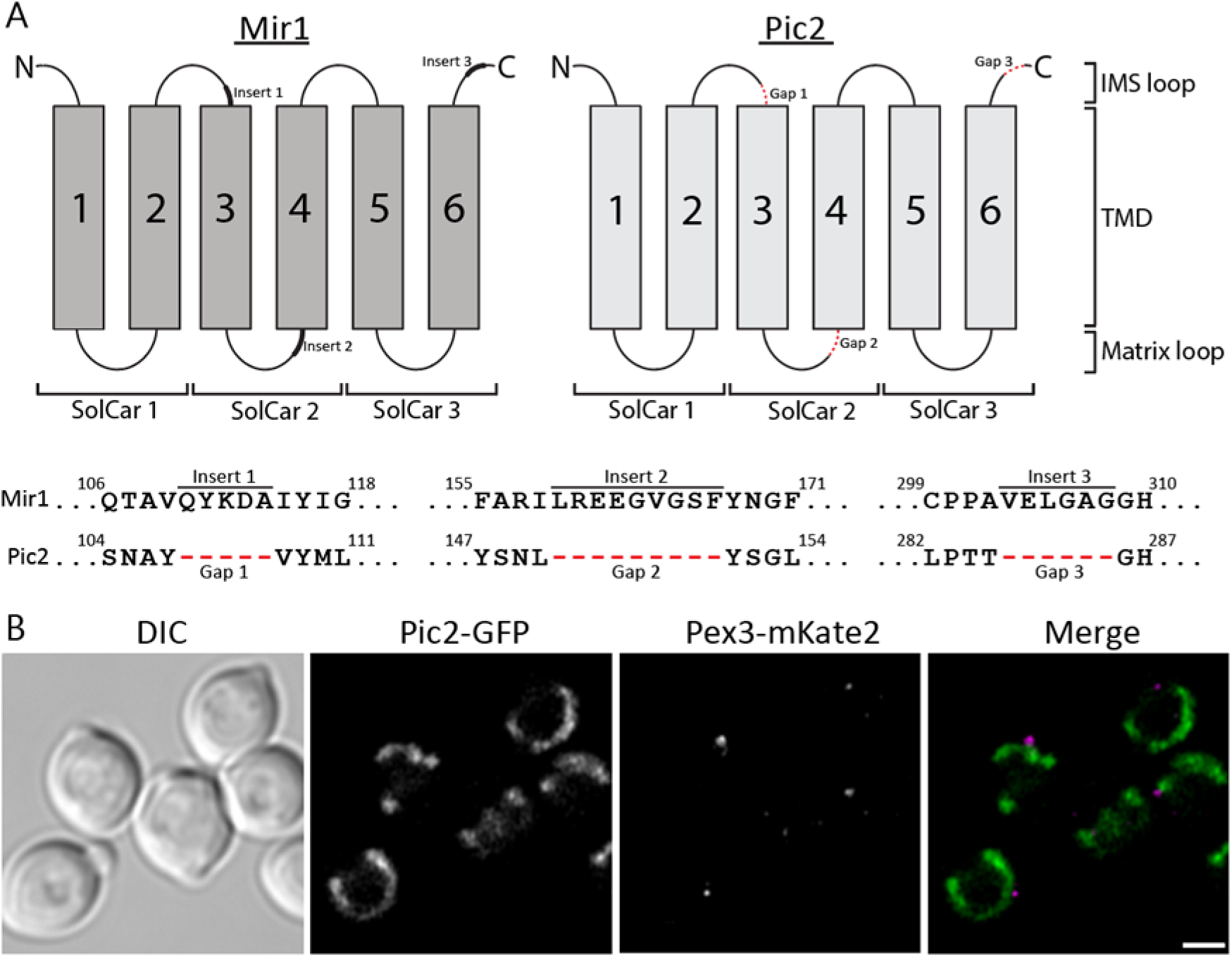
The Mir1 homolog Pic2 does not show a dual localization. **A)** Visual representation of the orientation Mir1 and Pic2 in MIM with numbered TMD (TMD 1-6), solute carrier domain (SolCar 1-3), the flexible loops facing the intermembrane space between each TMD (IMS loops) and the flexible loops facing the mitochondrial matrix between each TMD (Matrix loops). Red dashes in the flexible loop region of Pic2 indicate the topological location of the gap stretches (Gap 1-3), and at the corresponding topological locations on Mir1 are marked as insert (insert1-3) as bold line stretches. Pair wise sequence alignment of Mir1 and Pic2 visualizing the Mir1 inserts (underlined as Insert1-3) and the Pic2 gaps (marked in red dashed lines). Corresponding amino acids flanking the insert/gap sequences are numbered. **B)** Single focal plane confocal laser scanning microscopy Airyscan images of glucose-grown cells expressing C-terminal fusion of Pic2-GFP, under control of the endogenous promoters. Pex3-mKate2 was introduced as peroxisomal marker. The arrows indicate colocalization with Pex3- mKate2. Scalebar 1µm.

### The full-length Mir1 protein is required for targeting to peroxisomes

Because three gaps occur in Pic2 and Pic2 does not localize to peroxisomes, we reasoned that peroxisomal sorting information in Mir1 may occur in the three corresponding inserts in Mir1 (Fig. 5A).

To test this we generated Mir1 truncations containing GFP at the C-terminus (Fig. 6A). These fusion constructs were produced in a WT background, which contains the full length Mir1, because Δ*mir1* cells have a severe growth defect (Fig. S2). Mir1 is a member of the mitochondrial carrier family (MCF). Proteins of this family are typically composed of three repeated solute carrier (SolCar) domains each consisting of two transmembrane domains (TMDs) connected by flexible loops (Rampelt et al., 2020). To determine whether the individual Mir1 SolCar domains could target truncated Mir1 to peroxisomes, we generated Mir1 TMD1-2, TMD3-4 (containing insert 1 and 2), and TMD5-6 (containing insert 3) (Fig. 5A). TMD1-2 localized to tubular mitochondrial structures, but no co-localization was detected with Pex3-mKate2 (Fig. 6A). Unfortunately, it was not possible to determine the localization of TMD3-4 and TMD5-6 due to low GFP signal intensities (Fig. 6A). Based on this experiment, we therefore could not establish whether the three inserts play a role in peroxisomal sorting.

**Figure 6:**
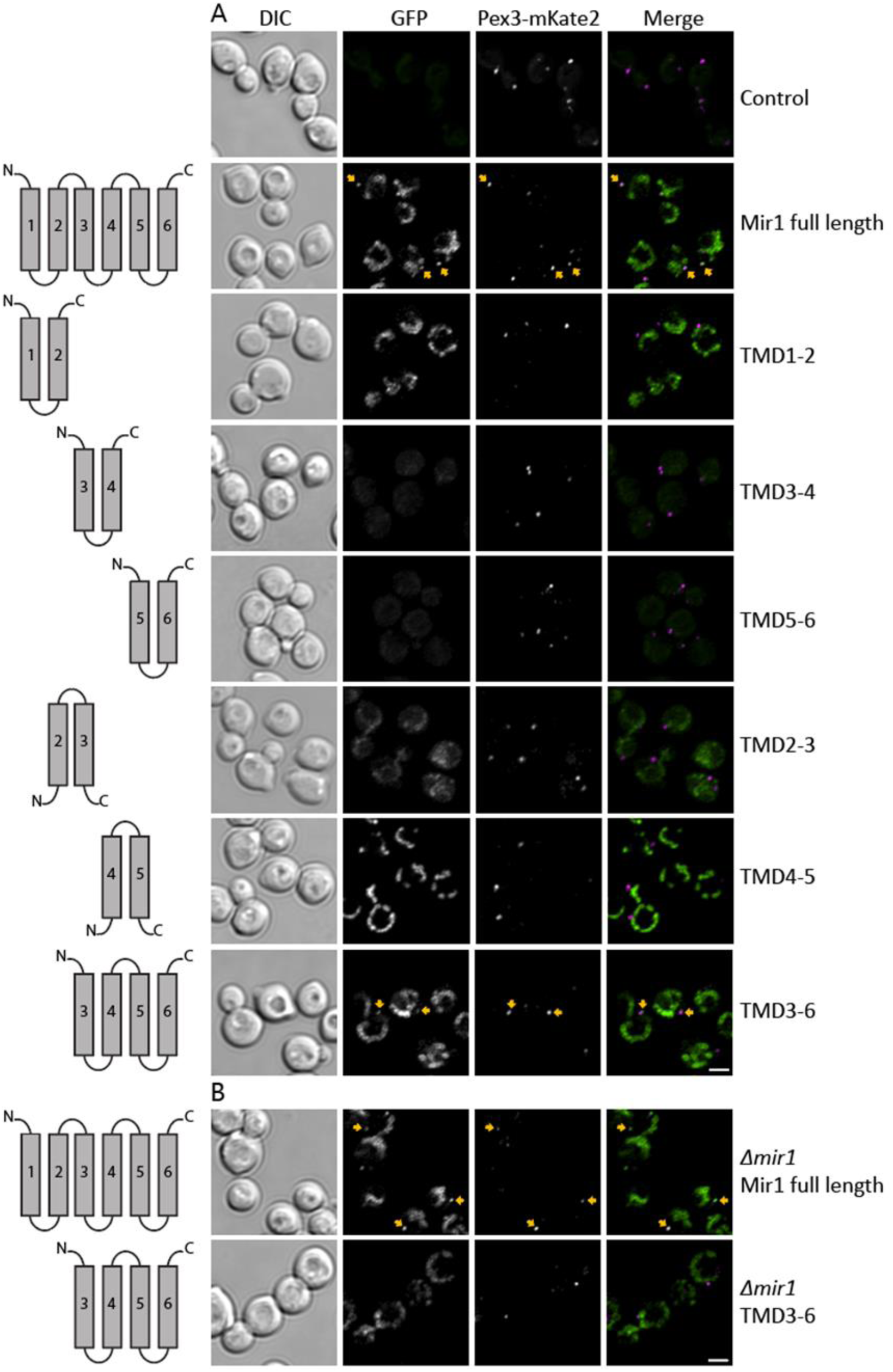
Localization studies of truncated Mir1 variants in WT and *Δmir1*. Single focal plane confocal laser scanning microscopy Airyscan images of glucose-grown WT cells overexpressing C-terminal fusions of **A)** full length Mir1 and truncated variants of Mir1 cloned into the genome also containing the full length untagged *MIR1* gene under control of its endogenous promoter. **B)** Images of glucose-grown *Δmir1* cells overexpressing C-terminal fusion of full length Mir1-GFP and the truncated version consisting of TMD3-6 cloned into genome of a *Δmir1* strain. Pex3-mKate2 peroxisomal marker. Scale bars 1µm.

We next tested other truncations, namely truncation TMD2-3 (containing insert 1) and TMD4-5 (containing insert 2). Both constructs showed mitochondrial structures, but no structures co-localizing with Pex3-mKate2 were observed (Fig. 6A). Finally, we analysed truncation TMD3-6 which contains all three insert. Interestingly, this construct showed again a dual localization on peroxisomes and mitochondria (Fig. 6A). This observation suggests that all three inserts are required for peroxisomal sorting.

To test whether TMD3-6 sorting depends on full length endogenous Mir1, we also expressed this construct in a Δ*mir1* strain. In Δ*mir1* cells the colocalization of Mir1 TMD3-6 with the peroxisomal marker was lost, while the full length Mir1-GFP control still showed the dual localization (Fig. 6B). Based on these observations, we conclude that only full length Mir1 is capable to sort to peroxisomes.

## Discussion

Many peroxisomal membrane transporters have been predicted based on metabolic pathways that are catalysed by peroxisomal enzymes, but so far only a few have been identified. It has been argued that additional transporters may not exist because the peroxisomal membrane contains pore forming proteins. Another explanation is that additional transporters do exist but have been overlooked because these are in fact mitochondrial transporters that dually localize to peroxisomes. Here we show that in the yeast *H. polymorpha* two mitochondrial inner-membrane transporters Aac2 and Mir1, not only localize to mitochondria, but also are peroxisomal.

An increasing number of proteins are identified to dually localize on mitochondria and peroxisomes. Initially, this mainly included peroxisomal matrix enzymes (e.g., Cat2 or Ptc5, (Elgersma et al., 1995) (Stehlik et al., 2020) (Bittner et al., 2022)and tail anchored proteins, which localize to the mitochondrial outer membrane and the peroxisomal membrane (e.g., yeast Fis1 and Gem1 (Cichocki et al., 2018).

Mitochondrial proteins are commonly detected in purified peroxisomal fractions, but these are generally considered to be contaminations as a result of the isolation procedure. In addition, peroxisomes are in physical contact with many other organelles *in vivo* at membrane contact sites, which may contributes to physical association of mitochondrial membranes to peroxisomes (Wu et al., 2023).

We observed that four mitochondrial transport proteins were specifically enriched in PMEM, namely *H. polymorpha* Mir1, Aac2, Por1 and Odc1. Interestingly, *S. cerevisiae* Mir1, Odc1 and Por1, and Aac2 (Pet9), were also detected in enriched peroxisomal fractions from *S. cerevisiae*, but considered to be contaminations (Marelli et al., 2004). Por1 is a very abundant mitochondrial outer membrane protein. We tried to localize both N- and C-terminal GFP fusions of Por1, but both were mislocalized to the cytosol. We also performed immune-electron microscopy using antibodies raised against *Sc*Por1. Even though these antibodies cross-reacted with *Hp*Por1, we could not detect specific labelling on mitochondria or peroxisomes. Therefore, we still cannot exclude that Por1 also has a dual localization on peroxisomes and mitochondria.

*S. cerevisiae* Odc1 and Odc2 have previously been implicated in linking mitochondrial and peroxisomal metabolism via the beta-oxidation when cells are cultivated on oleic acid (Tibbetts et al., 2002). We were unable to detect *Hp*Odc1 on peroxisomes by fluorescence microscopy. However, we cannot exclude that fluorescence intensities on peroxisomes are below the limit of detection. We did detect *Hp*Aac2 on mitochondria and peroxisomes. This was only observed in glucose-grown cells. We assume that Aac2 is also present on peroxisomes in methanol-grown cells, but likely below the limit of detection at these growth condition. Our observation is in line with the dual localization of *S. cerevisiae* Aac2 on mitochondria and peroxisomes (van Roermund et al., 2021). We also observed a dual localization of *Hp*Mir1, which like Aac2, is a carrier of the mitochondrial inner membrane. The localization of Mir1-GFP on peroxisomes was detectable both in glucose and methanol-grown cells.

Multiple processes generate phosphate in the peroxisomal matrix. In *H. polymorpha*, the Lon type protease Pln produces inorganic phosphate (Pi) through ATP hydrolysis (Aksam et al., 2007). This would indicate that Mir1 may export phosphate from the peroxisomal matrix. In *S. cerevisiae*, phosphate is generated in the peroxisomal matrix through degradation of CoA by Nudix hydrolase Pcd1 (Cartwright et al., 2000), activation of short -and medium chain fatty acids through ATP hydrolysis by Faa2 (Hettema et al., 1996) and dephosphorylation processes by Ptc5 (Stehlik et al., 2020).

*S. cerevisiae* Ant1 and its *H. polymorpha* homologue Pmp47 are implicated in ATP transport into peroxisomes. Recent studies in *S. cerevisiae* showed that ScAac2 also functions in ATP transport across the peroxisomal membrane (van Roermund et al., 2021). Based on these findings, we propose a new model for ATP and Pi transport across the peroxisomal membrane in *H. polymorpha* (Fig. 7). In this model, Mir1 is responsible for the export of Pi produced in the peroxisomal matrix by ATPases and phosphates. *H. polymorpha* only contains one Aac protein, which we called Aac2, because it is most homologous to ScAac2.

**Fig. 7:**
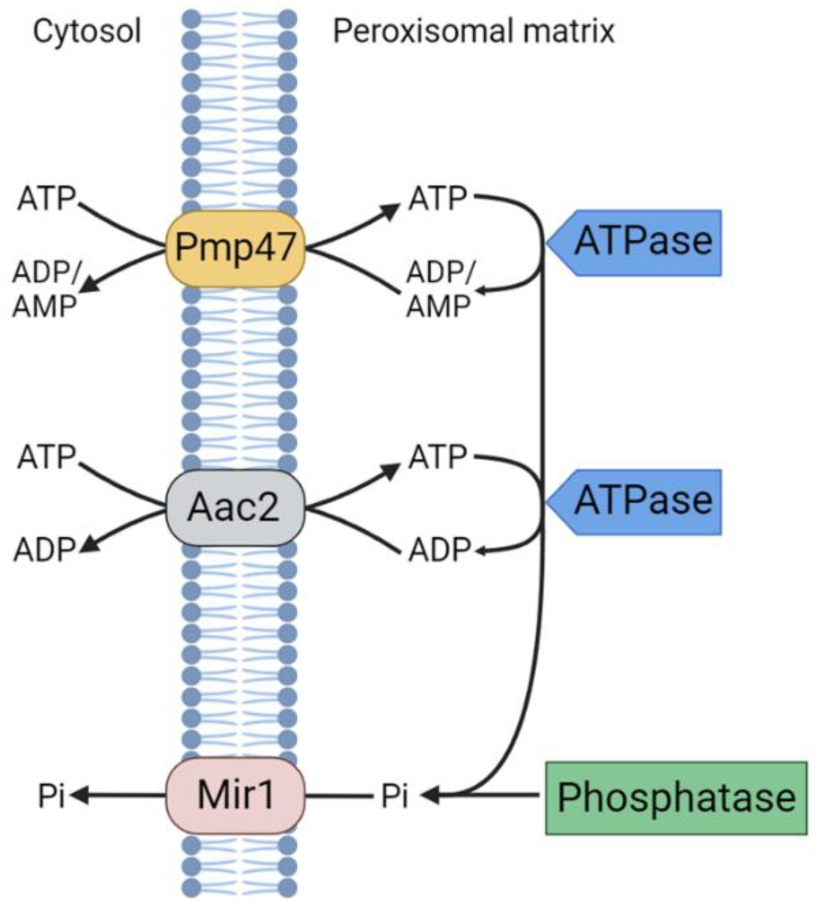
Proposed model for ATP:ADP(/AMP) exchange and Pi export across the peroxisomal membrane. Pmp47: ATP:ADP exchange. Aac2: ATP:ADP/AMP exchange. Mir1: Pi export. Peroxisomal ATPase (blue box) and phosphatase (green box) hydrolyse ATP hydrolyse and generates Pi.

S. *cerevisiae* Pex11 is a pore forming protein that is capable to allow diffusion of small solutes with a molecular mass below 300-400 Da (Mindthoff et al., 2016). Similarly, *in vivo* experiments in *S. cerevisiae* showed that small molecules could freely pass the peroxisomal membrane however with limitation for certain small hydrophobic molecules with a 300-400 Da cutoff (DeLoache et al., 2016). However, since the molecular weight of phosphate is 94,97 Da this molecule should be able to freely diffuse through these pores. Possibly, the Pex11 pore cannot accommodate phosphate. Alternatively, Mir1 is important when Pex11 is in a closed state.

Mitochondrial MCFs are known to contain multiple regions with mitochondrial targeting information (Horten et al., 2020; Kreimendahl et al., 2020). Similarly, the human peroxisomal MCF Pmp34 and the *Candida boidinii* peroxisomal MCF Pmp47 contain multiple regions with peroxisomal sorting information (mPTSs) (Jones et al., 2001; Wang et al., 2004). The majority of the Mir1 truncations which we analysed localized to mitochondria, indicating that Mir1 also contains multiple regions with mitochondrial sorting information. However, only full length Mir1 localized to peroxisomes.

Previously, the mPTSs for both human Pmp34 and *C. boidinii* Pmp47 were show to locate in the loops between TMD4-5 (Dyer et al., 1996; Honsho and Fujiki, 2001). However, we here show that a construct consisting of Mir1 TMD4-5 did not colocalize with the peroxisomal marker.

The Mir1 homolog, Pic2, did not localize to peroxisomes despite their high sequence identify. This corresponds to our observation that Pic2, which was detected in the PNS (Table S2) was not enriched in PMEM (Table S1). The only Mir1 truncation that exhibited peroxisomal colocalization in wild-type cells was Mir1 TMD3-6, which however lost its peroxisomal localization in *Δmir1*. Mir1 TMD3-6 contained all three inserts that are missing in Pic2. Possibly the three inserts are important to associate to full length Mir1, since in *Δmir1* Mir1 TMD3-6 is no longer peroxisomal.

In addition to the mitochondrial transporters, proteins from other organelles were enriched in the PMEM. For instance, the vacuolar protein Vac8 was also detected by MS in *S. cerevisiae* peroxisomal fractions (Marelli et al., 2004) and in complexes of the peroxisome-ER contact site protein *Sc*Pex30 (David et al., 2013). It is still not clear why Vac8 is often detected by proteomics of peroxisomal fractions. Studies on the role of *Hp*Vac8 failed to show a function of this protein in peroxisome biology (Singh et al., 2020).

Interestingly, we detected the plasma-membrane protein Pma2 in PMEM, which was also detected in purified peroxisomal fractions of *S. cerevisiae* (Marelli et al., 2004). Whether this protein plays a role in peroxisome biology still needs to be established. There was only one uncharacterized protein among the top 40 PMEM proteins, W1QB82. This protein contains an SAP domain, which binds DNA/RNA. Peroxisomes do not contain DNA or RNA, therefore we decided not to further analyse this protein.

Summarizing, proteomic analysis of peroxisomal membrane fractions combined with detailed fluorescence microscopy studies can yield interesting novel examples of dual localization. Moreover, the missing predicted peroxisomal transporters may all be proteins from other organelles that also localize to peroxisomes. Unfortunately, dual localization makes functional studies very hard. Since the absence of Mir1 has a severe effect on mitochondrial function and results in major growth defects, the *in vivo* analysis of the function of Mir1 in the peroxisomal membranes is very difficult.

## Materials and methods

### Strains and growth conditions

All *H. polymorpha* strains used in this study are listed in Table S3. Yeast cells were cultivated at 37°C in mineral medium (van Dijken et al., 1976) supplemented with 0.5% glucose, 0,5% methanol or a mixture of 0,25% methanol and 0,25% glucose. Leucine was added to a final concentration of 60 μg/ml, when necessary. Selection of positive transformants was performed on YPD plates (1% Yeast extract, 1% Peptone, and 1% Dextrose) containing 100 μg/ml zeocin (Invitrogen) and/or nourseothricin (Werner Bioagents). *Escherichia coli* DH5α were used for cloning.

### Plasmids and molecular techniques

Genes were expressed under control of the *TEF1*, *ADH1* or endogenous promoter (Table S4). For the expression under control of the *TEF1* and *ADH1* promoter the full-length gene was cloned.

All plasmids were linearized and integrated in the genome as described before (Faber et al., 1994; Saraya et al., 2012). All deletions were confirmed by Southern blotting. Plasmids used are listed in Table S4. DNA restriction enzymes were used as recommended by the suppliers (Thermo Scientific or New England Biolabs). Polymerase chain reactions (PCR) for cloning were carried out with Phusion High-Fidelity DNA Polymerase (Thermo Scientific). Colony PCR was carried out using Phire polymerase (Thermo Scientific). For DNA sequence analysis, the Clone Manager 5 program (Scientific and Educational Software, Durham, NC) was used. For FM localization studies, Pex3-mKate2 and Pex14-mCherry was used as peroxisomal marker.

A 243 nucleotide 5’stretch of the mitochondrial targeting sequence of *Neurospora crassa* F_0_ ATPase subunit 9 (Su9) (Westermann and Neupert, 2000), containing restriction site *Hind*III, ATG start codon, a twelve-nucleotide linker region and ending with a *Bgl*II restriction site was synthesized by GenScript into the pUC57 plasmid. By restriction digestion with *Hin*dIII and *Bgl*II, the Su9 fragment was ligated into the pHIPZ -mGFP plasmid to be C-terminally fused to mGFP. The fragment Su9-GFP from pHIPZ Su9-GFP was isolated upon HindIII and XbaI digestion and ligated into pHIPN18 mGFP using the same restriction enzymes.

Plasmid pHIPZ7 Mir1-GFP was constructed as follows: full length *MIR1* PCR fragment (930 bp) was amplified with primers L03-fw and L03-rev using genomic DNA of *H. polymorpha* yku80 as template and digested with restriction enzyme HindIII and BglII and ligated in HindIII and BglII digested plasmid pHIPZ7 Inp1-GFP (Krikken et al., 2020), producing pHIPZ7 Mir1-GFP. Subsequently, MunI-linearized pHIPZ7 Mir1-GFP was integrated into the genome of *H. polymorpha* yku80, *H. polymorpha* yku80 producing P*_PEX3_* Pex3-mKate2, *H. polymorpha Δmir1* and *H. polymorpha Δmir1* producing P*_PEX3_* Pex3-mKate2. Plasmid pHIPN Pex3-mKate2 was StuI-linearized and integrated into the genome of strain *H. polymorpha Δmir1*.

To create plasmid pHIPZ7 GFP-Mir1 PCR was performed on genomic DNA of *H. polymorpha* yku80 as template using primers 309.FW.eGFP.N-term. and 310.RV.eGFP.N-term. to obtain a 751 bp GFP fragment capable of C-terminal fusion to Mir1. Similarly, primers 311.FW.Mir1.N-term and 312.RV.Mir1.N-term was used to obtain a 967 bp Mir1 capable of N-terminal GFP fusion. The GFP fragment was HindIII and and BglII digested, and the Mir1 fragment was BglII and SalI digested. These two fragments were simultaneously ligated into HindIII and SalI digested pHIPZ7 Mir1-GFP backbone to create the pHIPZ7 GFP-Mir1 plasmid. MunI-linearized pHIPZ7 GFP-Mir1 was integrated into the genome of *H. polymorpha* yku80 and *H. polymorpha* yku80 producing P*_PEX3_* Pex3-mKate2.

To construct plasmid pHIPZ Mir1-GFP, PCR was performed on genomic DNA of *H. polymorpha* yku80 as template using primers Mir1-fw and Mir1-rev to produce a 773bp fragment and digested with restriction enzyme HindIII and BglII and ligated in HindIII and BglII digested plasmid pHIPZ -GFP, producing pHIPZ Mir1-GFP. Subsequently, EcoRI-linearized pHIPZ Mir1-GFP was integrated into the genome of *H. polymorpha* yku80, *H. polymorpha* yku80 producing P*_PEX3_* Pex3-mKate2, *H. polymorpha* NCYC495 producing P*_PEX14_* Pex14-mCherry, *H. polymorpha* NCYC495 *pex3 atg1* leu1.1, *H. polymorpha* NCYC495 *pex3 atg1* leu1.1 strain producing P*_PEX14_* Pex14-mCherry, *H. polymorpha* NCYC495 *pex19 atg1* leu1.1 and *H. polymorpha* NCYC495 *pex19 atg1* leu1.1 strain producing P*_PEX14_* Pex14-mCherry.

Genomic DNA of *H. polymorpha* yku80 was used as a template to generate a 882bp full length PCR fragment of Por1 using primers 292.FW.Por1.integr. and 293.RV.Por1.integr. The fragment was digested with restriction enzymes AfeI and BglII and ligated into a AfeI and BglII digested pHIPZ18 - GFP plasmid creating plasmid pHIPZ18 Por1-GFP. EcoRI-linearized pHIPZ18 Por1-GFP was integrated into the genome of strains including *H. polymorpha* yku80 and *H. polymorpha* yku80 producing P*_PEX3_* Pex3-mKate2. To construct a similar N-terminal GFP fusion, primers 317.FW.Por1.N-term. And 318.RV.Por1 produced a 883bp full length PCR fragment of Por1. The 883 bp PCR fragment was digested with restriction enzymes BglII and SalI and ligated in BglII and SalI digested plasmid pHIPZ7 GFP-Mir1 to produce plasmid pHIPZ7 GFP-Por1. MunI-linearized pHIPZ7 GFP-Por1 was integrated into the genome of *H. polymorpha* yku80 and *H. polymorpha* yku80 producing P*_PEX3_* Pex3-mKate2.

To construct plasmids containing truncations of Mir1, genomic DNA of *H. polymorpha* yku80 was used as a template to generate PCR fragments of: Mir1 TMD1-2 (330 bp) using primers 319.FW.Mir1_TMD1-2 and 320.RV.Mir1_TMD1-2; to create fragment containing Mir1 TMD3-4 (321 bp) with primers 334.FW.Mir1_TMD3- and 322.RV.Mir1_TMD3-4; to create fragment containing Mir1 TMD5-6 (369 bp) using primers 335.FW.Mir1_TMD5- and 324.RV.Mir1_TMD5-6; to create fragment containing Mir1 TMD2-3 (327 bp) using primers 336.FW.Mir1_TMD2- and 333.RV.Mir1_TMD-3; to create fragment containing Mir1 TMD4-5 (339 bp) using primers 337.FW.Mir1_TMD4- and 328.RV.Mir1_TMD1-5; to create fragment containing Mir1 TMD3-6 (663 bp) using primers 334.FW.Mir1_TMD3- and 324.RV.Mir1_TMD5-6. All Mir1 TMD PCR fragments were digested with restriction enzymes HindIII and BglII and ligated in HindIII and BglII digested plasmid pHIPZ7 Mir1- GFP, producing pHIPZ7 Mir1 TMD -GFP fusion truncation versions of Mir1. Subsequently, MunI-linearized pHIPZ7 Mir1 TMD -GFP plasmids were integrated into the genome of *H. polymorpha* yku80, *H. polymorpha* yku80 producing P*_PEX3_* Pex3-mKate2, *H. polymorpha Δmir1* and *H. polymorpha* yku80 producing P*_PEX3_* Pex3-mKate2.

Plasmid pHIPZ7 Aac2-GFP was constructed as follows: full length *AAC2* PCR fragment (921 bp) was amplified from genomic DNA of *H. polymorpha* yku80 as template using primers 6U7-fw and 6U7-rev. The 921 bp fragment was digested using HindIII and BglII and ligated into the HindIII and BglII digested plasmid pHIPZ7 Inp1-GFP, resulting in plasmid pHIPZ7 Aac2-GFP. MunI-linearized pHIPZ7 Aac2-GFP was integrated into the genome of *H. polymorpha* yku80 and *H. polymorpha* yku80 producing P*_PEX3_* Pex3-mKate2.

To construct pHIPZ18 Odc1-GFP a full-length fragment of *ODC1* was amplified from genomic DNA of *H. polymorpha* yku80 as template using primers ODC1-fw and ODC1-rev producing a PCR fragment of 909 bp which was digested using HindIII and BamHi and ligated into the HindIII and BamHi digested pHIPZ18 -GFP, producing plasmid pHIPZ18 Odc1-GFP. AgeI-linearized pHIPZ18 Odc1-GFP was integrated into the genome of *H. polymorpha* yku80 and *H. polymorpha* yku80 producing P*_PEX3_* Pex3-mKate2.

Construction of pHIPZ Pic2-GFP: amplification from genomic DNA of *H. polymorpha* yku80 as template using primers PIC2-fw and PIC2-rev resulting in a 563 bp fragment, which was digested with HindIII and BglII and ligated into HindIII and BglII digested pHIPZ Mir1-GFP, resulting in pHIPZ Pic2-GFP. Bsu36I -linearized pHIPZ Pic2-GFP was integrated into the genome of *H. polymorpha* yku80 and *H. polymorpha* yku80 producing P*_PEX3_* Pex3-mKate2.

### *MIR1* deletion

A *H. polymorpha* yku80 *MIR1* deletion strain was constructed by replacing the *MIR1* genomic region with a hygromycin resistance gene. First a PCR fragment of the hygromycin resistance marker was amplified from plasmid pHIPH4 as a template with primers MIR1del-fw and MIR1del-rev which each contains *MIR1* flanking regions producing a 2032 bp fragment. The resulting *MIR1* deletion cassette was transformed into *H. polymorpha yku80* cells to obtain strain Δ*mir1*. Deletion was confirmed by colony PCR and southern blotting.

### Chemostat cultivation

For peroxisome isolations cells were grown in a carbon limited chemostat using an 1,5L Applikon bioreactor and control units. Air was filtered through a cotton filled glass tubing followed by two air filters (Pall Acro, #4400) in parallel. Stirring was permanently at 450 rpm. The temperature was set at 37 °C. The pH was controlled by addition of 0,5 M sodium hydroxide to the culture. The pH was kept constant at 5,7.

The *H. polymorpha yku*80 Su9mt-GFP strain was extensively pre-cultivated in batch cultures in mineral media containing 0,5% glucose prior to inoculation of the bioreactor (van Dijken et al., 1976). The bioreactor was inoculated to a starting OD of 0,1 in mineral media containing 0,25% glucose. After 24 hours of batch cultivation the medium inlet was turned on. The feed consisted of mineral media supplemented with a mixture of 0,25% methanol and 0,25% glucose. Antifoam (Sigma Y-30 #A5758) was added to the media at a concentration of 7,5 ppm. The chemostat working volume was set to 1335 ml with a continuous dilution rate of D = 0,1h^-1^. After 5 working volume replacements 500 ml culture was harvested for peroxisome isolation.

### Peroxisome membrane enrichment

Preparation of post nuclear supernatant (PNS) was performed as previously described (van der Klei et al., 1998). PNS corresponding to 5-8 mg of protein was loaded onto a discontinuous sucrose gradient and centrifuged for 2,5h at 18.000rpm in a DuPont Instruments Sorvall SV-288 vertical rotor. After centrifugation 1 ml samples were harvested from the bottom. The fractions were analysed by western blotting using antibodies against Pex11 or GFP. Peroxisomal peak fractions were pooled. Fractions corresponding to 150 µg of protein were mixed 1:1 with Milli-Q water to dilute the high sucrose concentrations, followed by 1:1 mixing with ice-cold 0,1M Tris-HCl pH 8,0. After incubation for 15 min on ice, the sample was centrifuged at 200.000g at 4°C for 15 min. The supernatant was discarded and the pellet was resuspended in 1,5 ml 0,1M Na_2_CO_3_ pH 11,5, incubated on ice for 15 min followed by centrifugation at 200.000g at 4°C for 15 min (South and Gould, 1999). Membrane pellets of approximately 15 µg protein were stored at -20°C until mass spectrometry (MS) analysis. MS was performed on peroxisomal membrane fractions from 5 biological replicates as well as on the corresponding PNS fractions.

### Biochemical techniques

Sucrose density gradient fractions were first diluted 1:1 with Milli-Q water, prior to precipitation of the proteins with trichloroacetic acid (TCA) (McCammon et al., 1994). Protein concentrations were determined using Bio-Rad #5000006 protein assay. Sodium dodecyl sulphate–polyacrylamide gel electrophoresis (SDS-PAGE) using 12,5% discontinuous gels was performed as described previously (Baerends et al., 2000). Equal volumes of the sucrose gradient were loaded per lane.

Semi-dry transfer of proteins from the polyacrylamide gel to a nitrocellulose membrane (Amersham Protran 0.2 NC, #10600001) was performed as described previously ((Kyhse-Andersen, 1984). Blots were decorated with mouse monoclonal antiserum against GFP (Santa Cruz Biotechnology, Inc. #sc-9996) or rabbit polyclonal antisera against Pex11 (Krikken et al., 2009). Secondary goat anti-rabbit or goat anti-mouse antibodies conjugated to horseradish peroxidase (HRP) (Thermo Fisher Scientific, Invitrogen) were used for detection. Chemiluminescent (Amersham ECL Prime, #RPN2232) detection was performed according to the manufacturers guidelines (Bio-Rad, Chemidoc Imaging System). Blots were images using the Bio-Rad ChemiDoc imaging System.

### Fluorescence Microscopy (FM)

For Airyscan imaging, cells were fixed in 1% formaldehyde for 15 min on ice. Airyscan images were captured with a confocal laser scanning microscope (LSM800; Carl Zeiss) equipped with a 32-channel gallium arsenide phosphide photomultiplier tube (GaAsP-PMT), Zen 2009 software (Carl Zeiss), and a 63 × 1.40 NA objective (Carl Zeiss). The GFP signal was visualized by excitation with a 488 nm laser, and Mitotracker™, mCherry and mKate2 were visualized with a 561 nm laser. To stain mitochondria MitoTracker™ Red CMXRos (#M7512) was used according to manufactures protocol. ImageJ and Adobe Illustrator were used for image analysis.

### Correlative Light and Electron Microscopy (CLEM)

CLEM was performed for localization analysis as described previously (de Boer and van der Klei, 2023). 150-nm-thick cryo-sections were imaged with an Zeiss Axioscope A1 fluorescence microscope with a 63 × 1.40 NA objective (Carl Zeiss), CoolSNAP HQ2 digital camera and MICRO-MANAGER 1.4 software (Photometrics CoolSNAP HQ2, Birmingham, UK) were used for capturing images. The GFP signal was visualized with a 470/40 nm bandpass excitation filter, a 495 nm dichromatic mirror and a 525/50 nm band pass emission filter. The mKate2 fluorescence was visualized with a 587/25-nm bandpass excitation filter, a 605-nm dichromatic mirror, and a 647/70-nm bandpass emission filter. The DAPI signal was detected with a 380/30 bandpass excitation filter and a 460/50 nm bandpass emission filter. The grid was post-stained and embedded in a mixture containing 0.5% uranyl acetate and 1 % methylcellulose. A CM12 (Philips) transmission Electron microscope under 100 kV was used for the generation of double-tilt tomography series including a tilt range of 45° to −45° with 2.5° increments. To make CLEM images, FM and EM images were aligned using the eC-CLEM plugin (Paul-Gilloteaux et al., 2017) in Icy (http://icy.bioimageanalysis.org). The IMOD software package was used for reconstructing and 3 D rendering of the tomograms.

### Proteomics sample preparation

Protein levels were determined using discovery-based proteomics (using label free quantification) for relative protein concentrations (Wegrzyn et al., 2020). Briefly, in-gel digestion was performed on the PNS and enriched peroxisomal membranes using 300 ng trypsin (sequencing grade modified trypsin V5111; Promega) after reduction with 10 mmol/L dithiothreitol and alkylation with 55 mmol/L iodoacetamide proteins as described previously (Wolters et al., 2016).

### Discovery-based proteomics analyses

Discovery mass spectrometric analyses were performed on a quadrupole orbitrap mass spectrometer equipped with a nano-electrospray ion source (Orbitrap Q Exactive Plus, Thermo Scientific). Chromatographic separation of the peptides was performed by liquid chromatography (LC) on a nano-HPLC system (Ultimate 3000, Dionex) using a nano-LC column (Acclaim PepMapC100 C18, 75 µm x 50 cm, 2 µm, 100 Å, Dionex, buffer A: 0.1% v/v formic acid, dissolved in milliQ-H_2_O, buffer B: 0.1% v/v formic acid, dissolved in acetonitrile). 10-25 ug digested total protein starting material was injected using the µL-pickup method with buffer A as a transport liquid from a cooled autosampler (5 °C) and loaded onto a trap column (µPrecolumn cartridge, Acclaim PepMap100 C18, 5 µm, 100 Å, 300 µmx5 mm, Dionex). Peptides were separated on the nano-LC column using a linear gradient from 2-50% buffer B in 87 min at a flowrate of 300 nL/min. The mass spectrometer was operated in positive ion mode and data-dependent acquisition mode (DDA) using a top-15 method. MS spectra were acquired at a resolution of 70.000 at m/z 200 over a scan range of 300 to 1650 m/z with a AGC target of 3e^6^ ions and a maximum injection time of 50 ms. Peptide fragmentation was performed with higher energy collision dissociation (HCD) using a normalized collision energy (NCE) of 28. The intensity threshold for ions selection was set at 5.0 e^4^ with a charge exclusion of 1≤ and ≥6. The MS/MS spectra were acquired at a resolution of 17.500 at m/z 200, a AGC target of 5e^3^ ions and a maximum injection time of 50 ms and the isolation window set to 1.8 m/z. LC-MS raw data were processed with MaxQuant (version 1.5.5.1) (Cox and Mann, 2008).

Peptide and protein identification were carried out with Andromeda against the *H. polymorpha* Uniprot database (Identifier: #UP000008673. *Ogataea parapolymorpha* (strains ATCC 26012 / BCRC 20466 / JCM 22074 / NRRL Y-7560 / DL-1). Taxon ID: #871575) to which ATP9_NEUCR and GFP were added. Proteins were quantified with the MaxLFQ algorithm (Cox et al., 2014), including both razor and unique peptides and a minimum ratio count of two and filtered for the number of replicates. Finally, ranking tables using the median corrected log2-LFQ signal intensity from highest to lowest were made.

### Bioinformatic tools

Peptide sequence for *Hp*Mir1 (UniProt #W1QL03) and *Hp*Pic2 (UniProt #W1QKL5) were extracted from UniProt. The peptide sequences for W1QL03 and W1QKL5 were subjected to EMBOSS tool Pairwise Sequence Alignment for global alignment using the Needleman-Wunsch (EBLOSUM62, 10 gap penalty, 0.5 extend penalty) with Needle to reveal sequence identity, sequence similarity and sequence gap percentages (https://www.ebi.ac.uk/Tools/psa/emboss_needle/. Date used: 21-04-2022). Transmembrane domains were identified for W1QL03 and W1QKL5 using the DTU Health Tech bioinformatic service TMHMM 2.0 (https://services.healthtech.dtu.dk/services/TMHMM-2.0/. Date used: 21-04-2022). Solute carrier domains were identified for W1QL03 and W1QKL5 using the EMBL-EBI InterPro scan for Classification of protein families (https://www.ebi.ac.uk/interpro/search/sequence/. Date used: 21-04-2022).

Manual inspection of the top 40 log2-LFQ hits: Proteins from the *Ogataea parapolymorpha* DL-1 proteome (#UP000008673) without designated proteins names, were named after their closest homologs found in *Saccharomyces cerevisiae or Pichia pastoris*.

UniProt blastp of unnamed protein #W1QFR9 described as “ACB domain-containing protein” in the #UP000008673 proteome, revealed a high homology to *Pichia pastoris* Uniprot entry #C4R8D7 named “Autophagy-related protein 37” (Atg37; Nazarko et al., 2014, DOI: www.jcb.org/cgi/doi/10.1083/jcb.201307050). Thus, protein #W1QFR9 was given the description of “Autophagy-related protein 37” and “Atg37” as a protein name. Bioinformatic service https://www.uniprot.org/blast (date used 02-11-2023), with the BLOSUM62 matrix, E-Threshold 10, gapped “Yes”, HSps per hits “All” and was targeted against the “UniProtKB reference proteomes + Swiss-Prot” databases.

## Author contributions

MP and IK conceived the study. MP, JW, RB and AK performed the experiments. JW performed statistical analysis of proteomic data. MP wrote the original draft of the manuscript. All authors contributed to reviewing and editing the manuscript. MP and RB prepared the figures. All authors approved the submitted version.

## Acknowledgements

We thank you Robin Bakema for assisting in cloning and strain creation. This project has received funding from the European Union’s Horizon 2020 research and innovation programme under the Marie Skłodowska-Curie grant agreement No 812968.

## Supplementary figures

**Fig. S1:**
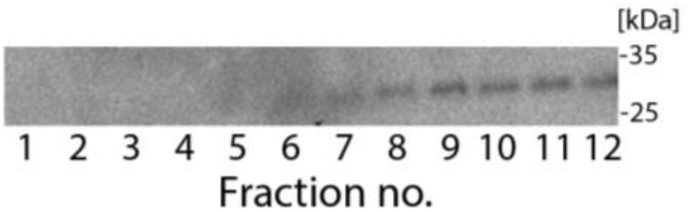
Long exposure of sucrose gradient fractions decorated with anti-GFP to detect the mitochondrial marker Su9-GFP. Western blot of sucrose gradient fractions 1-12 using anti-GFP antibodies. Equal volumes were loaded per lane. Long exposure (240 sec) of the same blot as shown in Fig. 1B anti-GFP (lower panel).

**Fig. S2:**
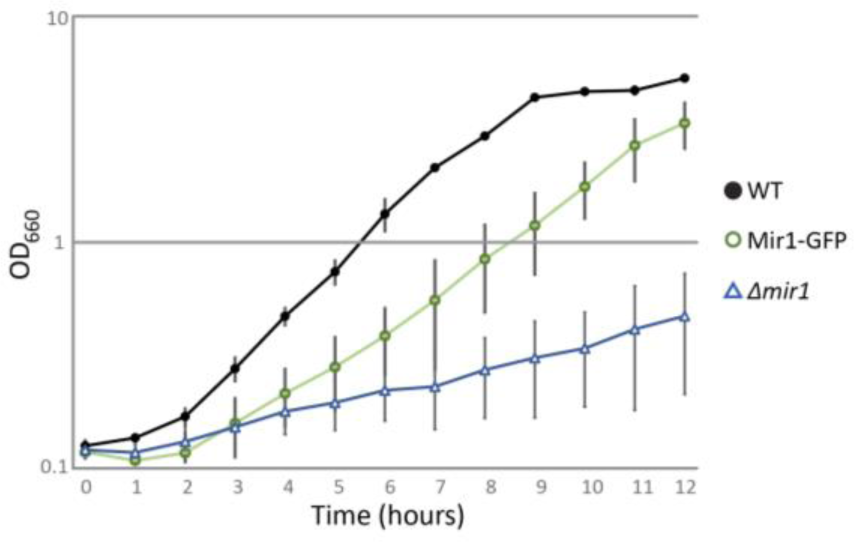
Growth curves of Mir1 strains. Growth curves of a *H. polymorpha* WT, *Δmir1* and a strain expressing Mir1-GFP under control of the endogenous promoter. Cells are grown on mineral medium containing glucose. The optical densities (y-axis) are expressed as absorbance at 660 nm (OD660) presented in logarithmic scale. Data are mean ± s.d. of two independent cultures.

## Supplementary Tables

**Table S1: Excel file with full table of proteins detected in PMEM, median corrected and ranked.**

**Table S2: Excell file with full table of proteins detected in PNS, median corrected and ranked.**

**Table S3:** Strains used for this study.

| Strain | Description | Reference |
| --- | --- | --- |
| Wild-type | *Hp.* NCYC495; leu1.1 | (Sudbery et al., 1988) |
| WT *yku80* | *Hp.* NCYC495, *YKU80::URA3*; leu1.1 | (Saraya et al., 2012) |
| WT:: $P_{PEX3}$ Pex3-mKate2 | *Hp. yku80* with integration of plasmid pHIPN Pex3-mKate2; $\text{Nat}^R$, *LEU2* | (Krikken et al., 2020) |
| WT:: $P_{ADH1}$ Su9mt-GFP | *Hp. yku80* with integration of plasmid pHIPN18 Su9mt-GFP | This study |
| WT:: $P_{MIR1}$ Mir1-GFP | *Hp. yku80* with integration of plasmid pHIPZ Mir1-GFP; $\text{Zeo}^R$, leu1.1 | This study |
| WT:: $P_{MIR1}$ Mir1-GFP::$P_{PEX3}$ Pex3-mKate2 | *Hp. yku80* $P_{PEX3}$ Pex3-mKate2 with integration of plasmid pHIPZ Mir1-GFP; $\text{Zeo}^R$, leu1.1 | This study |
| WT:: $P_{TEF1}$ Mir1-GFP | *Hp. yku80* with integration of plasmid pHIPZ7 Mir1-GFP; $\text{Zeo}^R$, leu1.1 | This study |
| WT:: $P_{TEF1}$ Mir1-GFP::$P_{PEX3}$ Pex3-mKate2 | *Hp. yku80* $P_{PEX3}$ Pex3-mKate2 with integration of plasmid pHIPZ7 Mir1-GFP; $\text{Zeo}^R$, leu1.1 | This study |
| WT:: $P_{TEF1}$ GFP-Mir1 | *Hp. yku80* with integration of plasmid pHIPZ7 GFP-Mir1; $\text{Zeo}^R$, leu1.1 | This study |
| WT:: $P_{TEF1}$ GFP-Mir1::$P_{PEX3}$ Pex3-mKate2 | *Hp. yku80* $P_{PEX3}$ Pex3-mKate2 with integration of plasmid pHIPZ7 GFP-Mir1; $\text{Zeo}^R$, leu1.1 | This study |
| WT:: $P_{TEF1}$ Aac2-GFP | *Hp. yku80* with integration of plasmid pHIPZ7 Aac2-GFP; $\text{Zeo}^R$, leu1.1 | This study |
| WT:: $P_{TEF1}$ Aac2-GFP::$P_{PEX3}$ Pex3-mKate2 | *Hp. yku80* $P_{PEX3}$ Pex3-mKate2 with integration of plasmid pHIPZ7 Aac2-GFP; $\text{Zeo}^R$, leu1.1 | This study |
| WT:: $P_{ADH1}$ Odc1-GFP | *Hp. yku80* with integration of plasmid pHIPZ7 Odc1-GFP; $\text{Zeo}^R$, leu1.1 | This study |
| WT:: $P_{ADH1}$ Odc1-GFP::$P_{PEX3}$ Pex3-mKate2 | *Hp. yku80* $P_{PEX3}$ Pex3-mKate2 with integration of plasmid pHIPZ7 Odc1-GFP; $\text{Zeo}^R$, leu1.1 | This study |
| WT:: $P_{ADH1}$ Por1-GFP | *Hp. yku80* with integration of plasmid pHIPZ7 Por1-GFP; $\text{Zeo}^R$, leu1.1 | This study |
| WT:: $P_{ADH1}$ Por1-GFP::$P_{PEX3}$ Pex3-mKate2 | *Hp. yku80* $P_{PEX3}$ Pex3-mKate2 with integration of plasmid pHIPZ7 Por1-GFP; $\text{Zeo}^R$, leu1.1 | This study |
| WT:: $P_{TEF1}$ GFP-Por1 | *Hp. yku80* with integration of plasmid pHIPZ7 GFP-Por1; $\text{Zeo}^R$, leu1.1 | This study |
| WT:: $P_{TEF1}$ GFP-Por1::$P_{PEX3}$ Pex3-mKate2 | *Hp. yku80* $P_{PEX3}$ Pex3-mKate2 with integration of plasmid pHIPZ7 GFP-Por1; $\text{Zeo}^R$, leu1.1 | This study |
| WT:: $P_{PIC2}$ Pic2-GFP | *Hp. yku80* with integration of plasmid pHIPZ Pic2-GFP; $\text{Zeo}^R$, leu1.1 | This study |
| WT:: $P_{PIC2}$ Pic2-GFP::$P_{PEX3}$ Pex3-mKate2 | *Hp. yku80* $P_{PEX3}$ Pex3-mKate2 with integration of plasmid pHIPZ Pic2-GFP; $\text{Zeo}^R$, leu1.1 | This study |
| $P_{PEX14}$ Pex14-mCherry | *Hp.* NCYC495 $P_{PEX14}$ Pex14-mCherry | (Knoops et al., 2014) |
| $P_{PEX14}$ Pex14-mCherry $P_{MIR1}$ Mir1-GFP | *Hp.* NCYC495 $P_{PEX14}$ Pex14-mCherry with integration of plasmid pHIPZ Mir1-GFP; $\text{Zeo}^R$, leu1.1 | This study |
| *pex3::atg1::leu1.1* | *Hp.* NCYC495 *pex3 atg1* leu1.1 | (Knoops et al., 2014) |
| *pex3::atg1::P_{PEX14}* Pex14-mCherry | *Hp.* NCYC495 *pex3 atg1* $P_{PEX14}$ Pex14-mCherry | (Knoops et al., 2014) |
| *pex3::atg1::P_{MIR1}* Mir1-GFP | *Hp.* NCYC495 *pex3 atg1* with integration of plasmid pHIPZ Mir1-GFP; $\text{Zeo}^R$, leu1.1 | This study |
| *pex3::atg1::P_{MIR1}* Mir1-GFP::$P_{PEX14}$ Pex14-mCherry | *Hp.* NCYC495 *pex3 atg1* $P_{PEX14}$ Pex14-mCherry with integration of plasmid pHIPZ Mir1-GFP; $\text{Zeo}^R$, leu1.1 | This study |
| *pex19:: atg1:: leu1.1* | *Hp. NCYC495 pex19 atg1 leu1.1* | (Knoops et al., 2014) |
| *pex19:: atg1:: P<sub>PEX14</sub> Pex14-mCherry* | *Hp. NCYC495 pex19 atg1 P<sub>PEX14</sub> Pex14-mCherry* | (Knoops et al., 2014) |
| *pex19:: atg1:: P<sub>MIR1</sub> Mir1-GFP* | *Hp. NCYC495 pex19 atg1* with integration of plasmid pHIPZ Mir1-GFP; Zeo<sup>R</sup>, leu1.1 | This study |
| *pex19:: atg1:: P<sub>MIR1</sub> Mir1-GFP::P<sub>PEX14</sub> Pex14-mCherry* | *Hp. NCYC495 pex19 atg1 P<sub>PEX14</sub> Pex14-mCherry* with integration of plasmid pHIPZ Mir1-GFP; Zeo<sup>R</sup>, leu1.1 | This study |
| WT:: *P<sub>TEF1</sub> Mir1_TMD1-2-GFP* | *Hp. yku80* with integration of plasmid pHIPZ7 Mir1_TMD1-2-GFP; Zeo<sup>R</sup>, leu1.1 | This study |
| WT:: *P<sub>TEF1</sub> Mir1_TMD1-2-GFP::P<sub>PEX3</sub> Pex3-mKate2* | *Hp. yku80 P<sub>PEX3</sub> Pex3-mKate2* with integration of plasmid pHIPZ7 Mir1_TMD1-2-GFP; Zeo<sup>R</sup>, leu1.1 | This study |
| WT:: *P<sub>TEF1</sub> Mir1_TMD3-4-GFP* | *Hp. yku80* with integration of plasmid pHIPZ7 Mir1_TMD3-4-GFP; Zeo<sup>R</sup>, leu1.1 | This study |
| WT:: *P<sub>TEF1</sub> Mir1_TMD3-4-GFP::P<sub>PEX3</sub> Pex3-mKate2* | *Hp. yku80 P<sub>PEX3</sub> Pex3-mKate2* with integration of plasmid pHIPZ7 Mir1_TMD3-4-GFP; Zeo<sup>R</sup>, leu1.1 | This study |
| WT:: *P<sub>TEF1</sub> Mir1_TMD5-6-GFP* | *Hp. yku80* with integration of plasmid pHIPZ7 Mir1_TMD5-6-GFP; Zeo<sup>R</sup>, leu1.1 | This study |
| WT:: *P<sub>TEF1</sub> Mir1_TMD5-6-GFP::P<sub>PEX3</sub> Pex3-mKate2* | *Hp. yku80 P<sub>PEX3</sub> Pex3-mKate2* with integration of plasmid pHIPZ7 Mir1_TMD5-6-GFP; Zeo<sup>R</sup>, leu1.1 | This study |
| WT:: *P<sub>TEF1</sub> Mir1_TMD2-3-GFP* | *Hp. yku80* with integration of plasmid pHIPZ7 Mir1_TMD2-3-GFP; Zeo<sup>R</sup>, leu1.1 | This study |
| WT:: *P<sub>TEF1</sub> Mir1_TMD2-3-GFP::P<sub>PEX3</sub> Pex3-mKate2* | *Hp. yku80 P<sub>PEX3</sub> Pex3-mKate2* with integration of plasmid pHIPZ7 Mir1_TMD2-3-GFP; Zeo<sup>R</sup>, leu1.1 | This study |
| WT:: *P<sub>TEF1</sub> Mir1_TMD3-6-GFP* | *Hp. yku80* with integration of plasmid pHIPZ7 Mir1_TMD3-6-GFP; Zeo<sup>R</sup>, leu1.1 | This study |
| WT:: *P<sub>TEF1</sub> Mir1_TMD3-6-GFP::P<sub>PEX3</sub> Pex3-mKate2* | *Hp. yku80 P<sub>PEX3</sub> Pex3-mKate2* with integration of plasmid pHIPZ7 Mir1_TMD3-6-GFP; Zeo<sup>R</sup>, leu1.1 | This study |
| WT:: *P<sub>TEF1</sub> Mir1_TMD4-5-GFP* | *Hp. yku80* with integration of plasmid pHIPZ7 Mir1_TMD4-5-GFP; Zeo<sup>R</sup>, leu1.1 | This study |
| WT:: *P<sub>TEF1</sub> Mir1_TMD4-5-GFP::P<sub>PEX3</sub> Pex3-mKate2* | *Hp. yku80 P<sub>PEX3</sub> Pex3-mKate2* with integration of plasmid pHIPZ7 Mir1_TMD4-5-GFP; Zeo<sup>R</sup>, leu1.1 | This study |
| *mir1::Hyg<sup>R</sup>* | *Hp. yku80 mir1 Hyg<sup>R</sup>*, leu1.1 | This study |
| *mir1 P<sub>TEF1</sub> Mir1-GFP* | *Hp. yku80 mir1* with integration of plasmid pHIPZ7 Mir1-GFP; Zeo<sup>R</sup>, leu1.1. Hyg<sup>R</sup> | This study |
| *mir1 P<sub>PEX3</sub> Pex3-mKate2* | *Hp. yku80 mir1* with integration of pHIPN Pex3-mKate2; Nat<sup>R</sup>, LEU2, Hyg<sup>R</sup> | This study |
| *mir1 P<sub>TEF1</sub> Mir1-GFP::P<sub>PEX3</sub> Pex3-mKate2* | *Hp. yku80 mir1 P<sub>PEX3</sub> Pex3-mKate2* with integration of plasmid pHIPZ7 Mir1-GFP; Zeo<sup>R</sup>, leu1.1. Hyg<sup>R</sup> | This study |
| *mir1 P<sub>TEF1</sub> Mir1_TMD3-6-GFP* | *Hp. yku80 mir1* with integration of plasmid pHIPZ7 Mir1_TMD3-6-GFP; Zeo<sup>R</sup>, leu1.1, Hyg<sup>R</sup> | This study |
| *mir1 P<sub>TEF1</sub> Mir1_TMD3-6-GFP::P<sub>PEX3</sub> Pex3-mKate2* | *Hp. yku80 mir1 P<sub>PEX3</sub> Pex3-mKate2* with integration of plasmid pHIPZ7 Mir1_TMD3-6-GFP; Zeo<sup>R</sup>, leu1.1, Hyg<sup>R</sup> | This study |

**Table S4:**
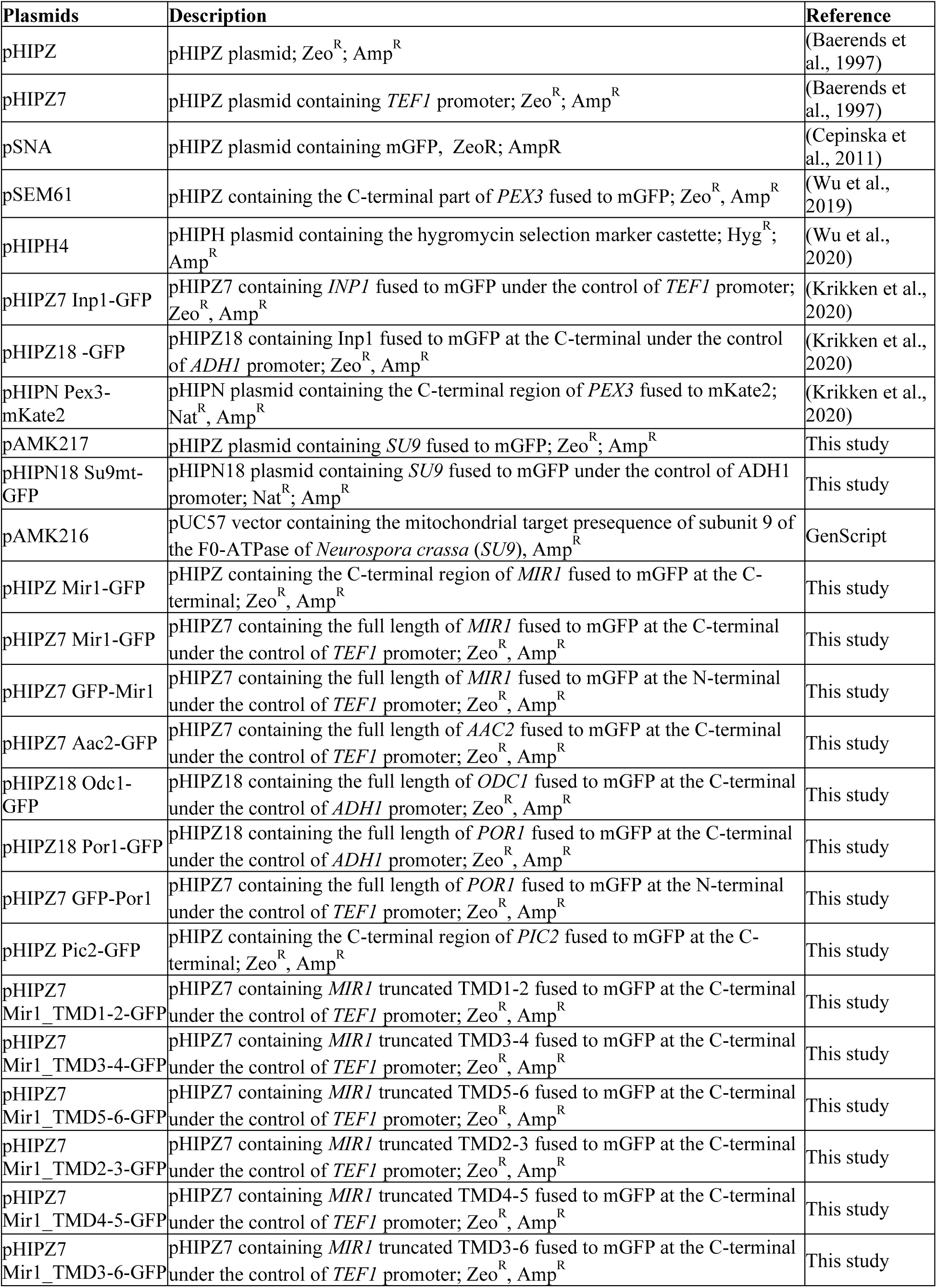
Plasmids used for this study.

**Table S5:**
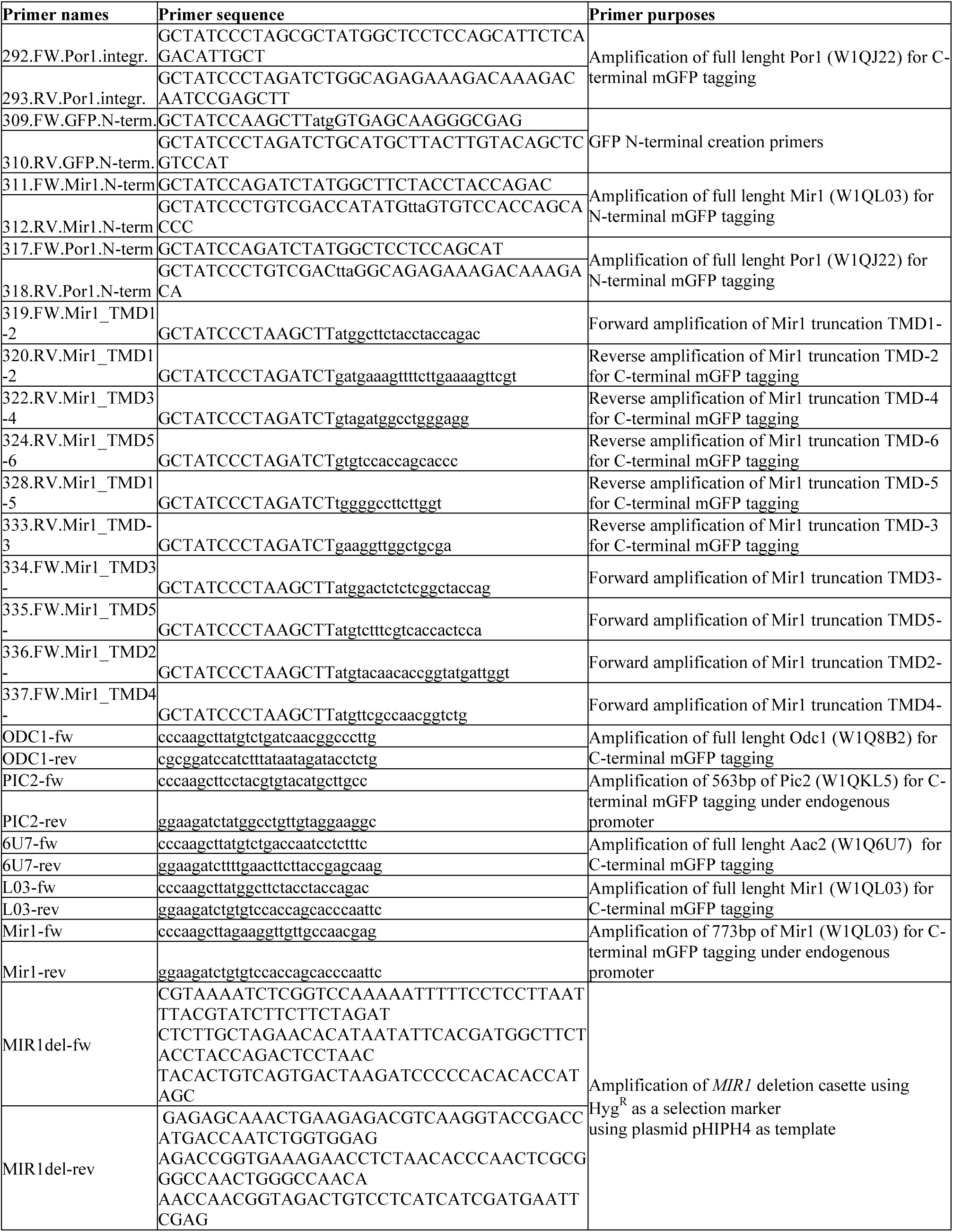
Primers used for this study.

